# A negative correlation between the rate coefficient of repair after photoinhibition of cold acclimated plants and the mean annual temperature of the habitats of *Arabidopsis thaliana* ecotypes

**DOI:** 10.1101/2024.07.07.602425

**Authors:** Riichi Oguchi, Soichiro Nagano, Ana Pfleger, Hiroshi Ozaki, Kouki Hikosaka, Barry Osmond, Wah Soon Chow

**Author notes:** Corresponding author: Dr. Riichi Oguchi.

## Abstract

Both the activity of photosynthesis and the repair of damaged photosystems decline in cold environments, which may increase the extent of the damage of photosynthetic machinery by light, namely photoinhibition. We hypothesized that plants in colder habitats may possess greater tolerance to photoinhibition, especially in low temperature conditions.

We measured the rate of photoinhibition, rate of photoinhibition repair and other thylakoid activities in cold environments using 298 *Arabidopsis thaliana* ecotypes and studied the relationships among the indicators of photoinhibition tolerance and climatic data of the habitat of each ecotype. The plants acclimated to cold conditions (12°C) for three days showed a negative correlation between the rate of photoinhibition repair at 5°C and the mean annual temperature of habitats, although we could not see this correlation with the control plants grown in 22°C. This result would indicate that the acclimation capacity of photoinhibition tolerance in cold conditions can affect the distribution of plants especially in colder regions.

## Introduction

Plants utilize sun light energy to drive photosynthesis, but strong sun light may cause damage to photosynthetic machineries and decrease the growth of plants, called photoinhibition (Jones & Kok, 1966; Osmond *et al*., 1997; Raven, 2011). Although the mechanisms of the photoinhibition are still controversial (Tyystjärvi, 2013; Nishiyama & Murata, 2014; Li *et al*., 2018; Kono *et al*., 2022), plants have many mechanisms to mitigate photoinhibition, such as chloroplast movement to avoid excessive light absorption (Senn, 1908; Kasahara *et al*., 2002), non-photochemical dissipation of excess energy from photosystems (Demmig *et al*., 1987; Niyogi *et al*., 1998), mechanisms of scavenging reactive oxygen species produced by excessively strong light energy (Nakano & Asada, 1981; Kojima *et al*., 2007; Khorobrykh *et al*., 2020), systems for repairing damaged photosystems (Aro *et al*., 1993; Kudoh & Sonoike, 2002; Kato *et al*., 2012), state-transitions to balance energy absorption of two photosystems (Bellafiore *et al*., 2005; Rochaix, 2014), consumption of excess light energy by cyclic electron flow (Munekage *et al*., 2004; Kono *et al*., 2014; Yamamoto & Shikanai, 2019), and so forth. The evolution of so many mechanisms of photoprotection indicates natural selection pressure for photoinhibition tolerance.

However, we still have insufficient information about the effect of photoinhibition on the distribution and diversification of plant species. This may be related to the characteristics of the variation of light intensity. Light intensity can strongly vary in space on small scales such as in the light gradients inside a canopy, and in time even in the same position such as the diurnal variation or seasonal variation, in the understory of deciduous forests (Valladares *et al*., 1997; Rothstein & Zak, 2001). Therefore, the variation of light intensity is not necessarily larger in the macro scale than in the micro scale geographically, which is contrast to temperature and precipitation. In consequence, few previous studies have studied the geographical variation of photoinhibition tolerance among species or ecotypes, although it has been well documented that sun plants have higher photoinhibition tolerance than shade plants (Öquist, 1992; Gomez-Aparicio *et al*., 2006). Nevertheless, the extent of photoinhibition strongly depends on environmental factors in addition to light intensity and increases in stressful conditions (Alves *et al*., 2002; Gururani *et al*., 2015). For example, diminished carboxylation activity at low temperature causes an imbalance between light energy absorption by photosystems and energy utilization by photosynthesis (Huner *et al*., 1998). The rates of repair of damaged photosystems also decrease at low temperature (Greer *et al*., 1986; Lee *et al*., 2001), which enhances photoinhibitory damage (see also Figure 1a). Therefore, the damage due to photoinhibition can be more severe in colder habitats at higher latitudes and higher altitudes (Miguez *et al*., 2015).

**Figure 1.**
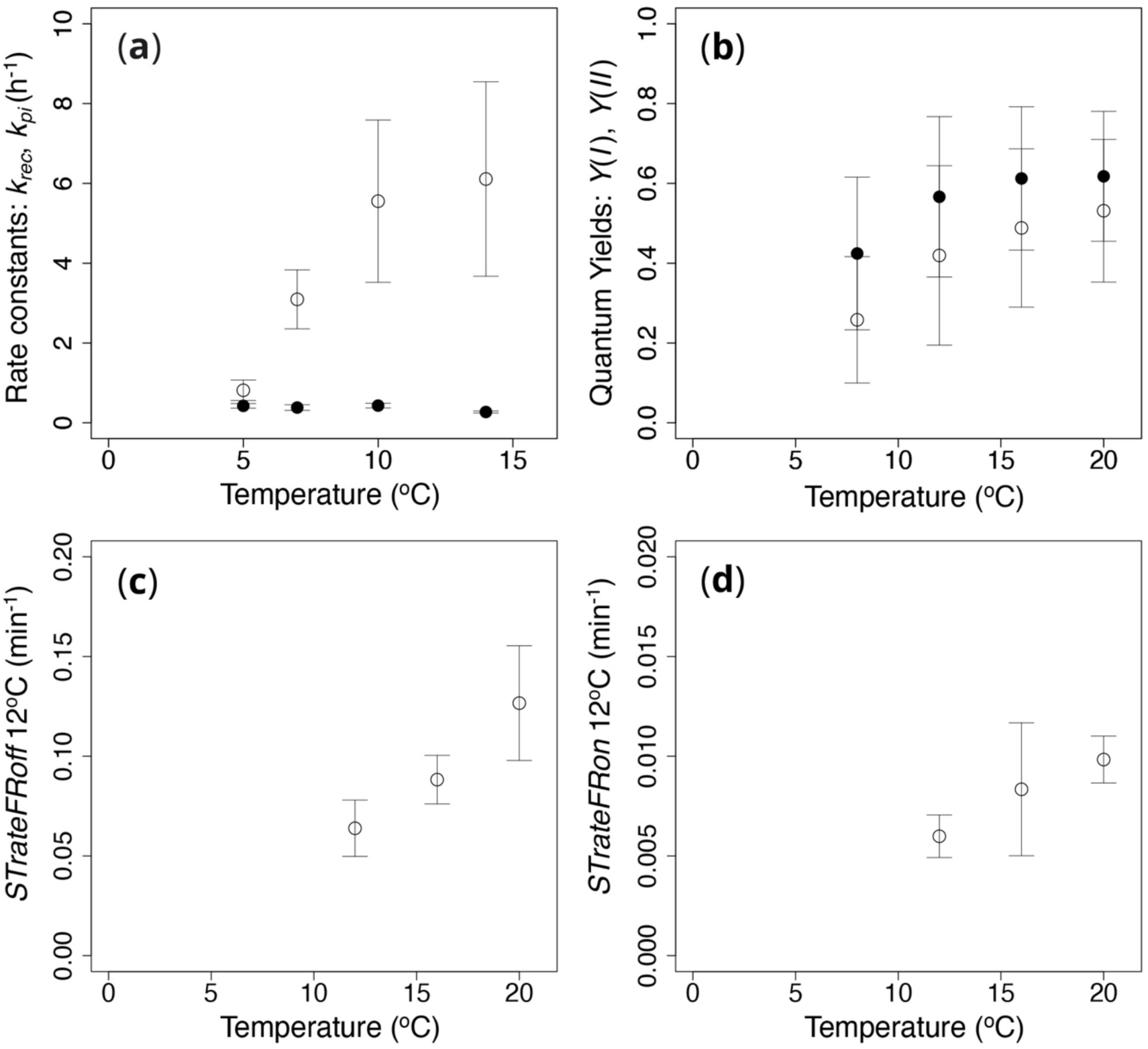
Temperature response of the rate coefficient of Photosystem II (PSII) photoinhibition (*k_pi_*, filled circles) and PSII photoinhibition repair (*k*_rec_, open circles) of Col-0 ecotype of *Arabidopsis thaliana* (a). The rate coefficients were calculated with the data of plants photoinhibited for 0, 30 and 60 minutes, with or without lincomycin (see text for the detail). PSII activity was estimated as the maximum quantum yield of PSII (*F_v_*/*F_m_*) using chlorophyll fluorescence. Temperature response of the quantum yield of photochemical energy conversion in PSII (*Y*(*II*): open circles in Panel b); the quantum yield of photochemical energy conversion in PSI (*Y*(*I*): closed circles in Panel b); the rate of state-transitions from State I to State II (*STrateFRoff*: Panel c); and the rate of state-transitions from State II to State I (*STrateFRon*: Panel d). The quantum yields of PSII and PSI and the rate of state transition were measured with chlorophyll fluorescence, P700 redox signal and light induced chlorophyll fluorescence transient (LIFT), respectively, using TFA08 ecotype of *Arabidopsis thaliana*.

We hypothesized that plants with higher photoinhibition tolerance in low temperature conditions have undergone selection in colder habitats. It has been reported that photoinhibition tolerance is greater in an ecotype of tomato (*Lycopersicon hirsutum*) from a higher altitude (Jung *et al*., 1998), and is greater in ecotypes of oak (*Quercus virginiana*) and Antarctic pearlwort (*Colobanthus quitensis*) from a higher latitude (Cavender-Bares, 2007; Bascunan-Godoy *et al*., 2012). These previous studies compared the extent of photoinhibition in a strong light condition using only two ecotypes; therefore, information about the relationships between habitat environments and photoinhibition tolerance, and about the relative contribution of various mechanisms of tolerance towards photoinhibition remains inadequate.

In the present research, we studied the variation of photoinhibition tolerance among 298 *Arabidopsis thaliana* ecotypes collected at various latitudes and altitudes all over the world, and sought correlations among the component mechanisms of tolerance to photoinhibition, leaf photosynthetic traits and environmental factors in original habitats. We specifically focused on the repair capacity of low-temperature-damaged Photosystem II, using plants that were either non-hardened or hardened by prior exposure to low temperature.

## Materials and Methods

### Plant Growth

*Arabidopsis thaliana* ecotypes were obtained from the Arabidopsis Biological Resource Center (ABRC). The list of used ecotypes is shown in the supporting information (Table S1). Plants were grown in a growth chamber in which the light intensity and temperature were controlled at 260 µmol m^−2^ s^−1^ and 22°C, respectively (10 h light and 14 h dark). For the photoinhibition experiments conducted in Japan, seeds were sown on rockwool blocks (3.6 × 3.6 × 4.0 cm size) and hydroponic culture solution (1000-times diluted Hyponex; N:P:K = 6:10:5, Hyponex-Japan, Osaka, Japan) was given 7 and 17 days after seed sowing. On 22 days after seed sowing, plants were transferred from the 22°C growth chamber to a smaller growth chamber, in which the light intensity and temperature were controlled at 18.2 µmol m^−2^ s^−1^ and at 12°C (for cold acclimation) or 22°C (for control), and grown for 3 days. Then, the first and the second leaves of the plants were harvested for photoinhibition treatments 25 days after seed sowing. For the measurements of photosystems activity and state-transitions conducted in Australia, seeds were sown on filter paper for the germination and after the germination seedlings were transferred to 6.65 × 6.65 × 9.5 cm pots with soils supplemented by a slow release fertilizer (‘Osmocote’, Scott Australia Pty. Ltd., Bella Vista, Australia). For the enhancement of seed germination, the seeds on filter paper were kept in the dark and cold (5°C) for four days before they were moved to a 22°C growth chamber. We used the plants, which rosette size (diameter) was larger than 5 cm, to obtain sufficient signals for measurements of photosystems activity and state-transitions. Therefore, the size of plants was larger than that of the photoinhibition experiments in Japan. The days needed for the sufficient growth described above varied depending on ecotypes and plants, which resulted in the variation of plant age (from 28 to 68 days after seed sowing).

### Photoinhibition measurement

The harvested leaves were immersed in to remove air bubbles of the abaxial surface and soon floated on 1 mM lincomycin solution as an inhibitor of photosystems repair, or water as the control. After 30 min of the lincomycin or water treatment in the dark, the leaves were photoinhibited by strong white LED light (1000 µmol m^−2^ s^−1^) at 5°C or 10°C. The duration of the photoinhibition treatment was 0, 30 and 60 min. After 30 min dark treatment to relax non-photochemical quenching, the maximum photochemical efficiency of photochemistry in photosystem II (PSII) was measured as *F_v_*/*F_m_* with a chlorophyll fluorescence imaging system (FluorCam, Photon Systems Instruments, Průmyslová, Drásov, Czech Republic). The rate coefficient of photoinhibition (*k_pi_*) and the rate of photoinhibition-repair (*k_rec_*) were calculated by fitting the relationship between photoinhibition time (*t*) and normalized *F_v_*/*F_m_* (denoted as *a* to represent the active fraction of PSII) with Equations 1 and 2, respectively as follows (Tyystjarvi *et al*., 1992; Wunschmann & Brand, 1992).

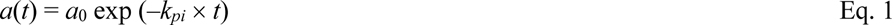

Equation 1 was fitted for the data obtained with lincomycin.

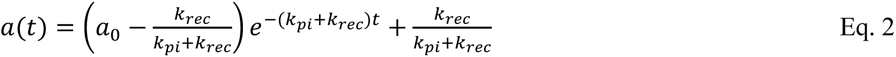

Equation 2 was fitted for the data obtained without lincomycin. In both equations, the initial *F_v_*/*F_m_* was normalized to give *a*_0_ = 1.

We also measured two indicators of energy-dependent non-photochemical quenching, *qE*, by measuring the non-relaxed chlorophyll fluorescence yield just after the photoinhibition treatment: (*F_v_*/*F_m_*)*_NR_*. The first indicator is the increment of *F_v_*/*F_m_* during the relaxation of non-photochemical quenching (*F_v_*/*F_m_* – (*F_v_*/*F_m_*)*_NR_*), which should indicate the strength of energy-dependent non-photochemical quenching. The second indicator is (*F_v_*/*F_m_* /(*F_v_*/*F_m_*)*_NR_* – 1), which we called *qE* indicator. This indicator should mean *k_E_* /(*k_F_* + *k_D_* + *k_P_*), where *k_E_*, *k_F_*, *k_D_* and *k_P_* indicate rate constants for the de-excitation of excited chlorophyll molecules by energy-dependent quenching, by fluorescence, by nonradiative decay process and by photochemistry, respectively (see Supporting Information for the detail). The indicators were calculated by using the data of leaf samples without lincomycin after 30 minutes photoinhibition treatment.

### Measurements of the activities of the photosystems

For the measurement of the activity of PSII, chlorophyll fluorescence yields were measured with a fluorometer (PAM101, 102 and 103; Walz, Effeltrich, Germany). The maximal (*F_m_*′), steady state (*F_s_*′) and minimal (*F_o_*′) fluorescence yields were determined under actinic light (600 μmol m^−2^ s^−1^) supplied by a halogen lamp (KL1500, Walz) at 12°C at saturating CO_2_. The quantum yield of PSII electron transport *Y*(*II*) was calculated as (*F_m_*′ – *F_s_*′)/ *F_m_*′ (Genty et al. 1989). For the measurement of the activity of PSI, immediately after the chlorophyll fluorescence measurements, the redox change of P700 was measured with a dual-wavelength (820/870 nm) unit (ED-P700DW) attached to a PAM fluorometer (Walz) and used in the reflectance mode (Chow & Hope, 2004). It was shown that the variation of the relative size of the P700 redox signal with far-red irradiance was practically identical for measurements in the reflectance or transmission modes (Kim *et al*., 2001). After the measurement of steady state P700 (*P_s_*′) under the same actinic light (600 μmol m^-2^ s^-1^), strong far-red light was given to deplete electrons from the inter-system chain and a saturation light pulse was applied to obtain maximally-oxidized P700 (*P_m_*′) (see Kou et al. (2013) for the detail). The quantum yield of PSI electron transport *Y*(*I*) was calculated as (*P_m_*′ – *P_s_*′)/*P_m_*, where *P_m_* was measured with the leaf sample illuminated by continuous weak far-red light to which was added a saturating pulse of white light (Klughammer & Schreiber, 1994; Siebke *et al*., 1997; Klughammer & Schreiber, 2008). The timing of the flash and the start of data acquisition, and the repetition rate of flash illumination, were controlled using a pulse/delay generator (Model 565 or 575; Berkeley Nucleonics Corporation, San Rafael, CA, USA). The analog output from the fluorometer was digitized and stored in a computer using a home-written program (by the late A. B. Hope).

### Measurement of state-transitions

Fluorescence transients in response to different light quality (Light I: white LED light mimicking sun light spectrum with added far-red light; and Light II: the same white LED light without far-red light, Artificial Sunlight Research Module GEN II, SL Holland, Breda, the Netherlands) were measured with a commercially available Light Induced Fluorescent Transient system (LIFT, Soliense Inc., Shoreham, NY, USA). The LIFT utilizes short-duration, high-frequency blue flashes (flashlets: 470 nm) to induce fluorescence changes in leaves caused by the increase in the proportion of closed reaction centers of PSII (Q_A_ saturation sequence). After the Q_A_ saturation sequence, LIFT also decreases the frequency of blue flashlets to allow reopening of the PSII reaction centers (Q_A_ relaxation sequence). The chlorophyll fluorescence from leaves was monitored in these two sequences to calculate the maximal (*F*_m_′), steady state (*F_s_*′) and minimal (*F_o_*′) fluorescence yields by fitting the Fast Repetition Rate (FRR) model (Kolber *et al*., 1998; Osmond *et al*., 2017).

The change of light quality from Light II to Light I gradually increases *F_m_*′ intensity measured by LIFT, which is caused presumably by the migration of a subpopulation of LHCII from PSI to PSII during state-transitions as previously validated using state transition mutants of *A*. *thaliana* (Osmond *et al*., 2019). The rate of the increase of *F_m_*′ intensity was used to calculate the rate of state-transition from State II to State I by a linear fitting (Figure S1). The change of light quality from Light I to Light II decreases *F_m_*′ intensity, the rate of which was used to calculate the rate of state-transition from State I to State II. Plants were put inside a temperature-controllable chamber having a transparent door, in which the air temperature was controlled to 12°C. The measurements by the LIFT were conducted with the blue flashlets and fluorescence passing through the transparent door.

### Measurement of leaf mass per area

We used the data of leaf mass per area (LMA) gained in our previous report (Oguchi *et al*., 2016), where 44 *Arabidopsis* ecotypes were grown in growth chambers (Biotron; NK system, Osaka, Japan). Three of six identical chambers were set to the ambient CO_2_ condition (380-400 μmol mol^−1^) and the other three chambers were set to the elevated CO_2_ condition (800 μmol mol^−1^). Growth temperature and humidity were set to 20°C and 60 %, respectively. The photoperiod was 10 h day^−1^ (130 μmol m^−2^ s^−1^ for a total of 9 h and 660 μmol m^−2^ s^−1^ for 1 h in the mid-photoperiod). The growth light source was metal-halide lamps. Plants were grown in pots (5 cm diameter and 5 cm height) filled with river sand. At 12 and 28 days after seed sowing, a hydroponic culture solution (Hewitt & Smith, 1975), in which the N concentration was reduced to half of the original concentration, was given in volumes of 20 and 40 ml per plant. At 38 days after the seed sowing, the largest leaf from each plant was harvested and scanned for measuring leaf area. After the desiccation in an oven (80°C), the dried mass of the largest leaf was measured for the calculation of leaf mass per area.

### Climate data

We used the climate data of a gridded climatology of 1961-1990 monthly means: CRU CL v. 2.0 (https://crudata.uea.ac.uk/cru/data/hrg/tmc/) provided by University of East Anglia (New *et al*., 2002). The data covered all land areas excluding Antarctica at 10’ resolution. The annual average of temperature, relative humidity and wind speed were calculated from the monthly average of the temperature, relative humidity and wind speed of the CRU data at the geographically closest point to the habitat latitude/longitude data of ecotypes, which were mainly obtained from ABRC. Altitudes were also obtained from the CRU data at the closest point to the habitat latitude/longitude data of ecotypes. The range of mean annual temperature, the lowest monthly temperature, and the highest monthly temperature of the habitat was –2.8 to 17.9°C, –17.3 to 13.9°C, and 5.2 to 30.2°C, respectively.

### Statistical analysis

The calculation, fitting and statistical analysis were conducted with R statistical software (version 4.2.2; R Foundation for Statistical Computing; available from http://www.R-proje ct.org).

## Results

Col-0 ecotype of *Arabidopsis thaliana* showed a strong temperature dependency in the repair rate coefficient after PSII photoinhibition (*k_rec_*), though its photoinhibition rate coefficient in the absence of repair (*k_pi_*) did not show temperature dependency (Figure 1a). According to these temperature dependencies, we have focused on the variations in *k_rec_* and *k_pi_* among ecotypes at 10 and 5°C, where *k_rec_* changed markedly between the two temperatures. We expected a correlation between *k_rec_* and the habitat temperature of each ecotype in these low temperature conditions, and conducted photoinhibition experiments using 189 and 298 *A*. *thaliana* ecotypes at 10 and 5°C, respectively. However, there was no statistically significant correlation between *k_rec_* and the habitat temperature of each ecotype both in 10 and 5°C (Figure 2a and b). A statistically significant correlation between the photoinhibition repair rate at 5°C and the habitat temperature was observed only after plants were acclimated to lower temperature (Figure 2b blue symbols; plants were grown at 22°C and acclimated to 12°C for 3 days). The low temperature acclimation significantly increased *k_rec_* at 5°C and the extent of the increase was larger in the ecotypes that originated from lower-temperature habitats. There was no statistically significant correlation between the habitat temperature and *k_pi_* at both 10 and 5°C (Figure 3a and b), but the cold-acclimated plants showed a significant correlation between them at 5°C (Figure 3b). The low temperature acclimation significantly decreased the photoinhibition rate coefficient at 5°C. We also tested multiple models of the relationships among *k_rec_* and environmental parameters (e.g. average temperature, altitude, average relative humidity and average wind speed of habitats) using the stepwise model selection of MASS package of R. The selected model with the smallest AIC (Akaike’s information criterion) was the relationship between *k_rec_* and average habitat temperature (Table 1). Multiple models of the relationships among *k_pi_* and environmental parameters were also tested as above. The selected model with the smallest AIC was the relationship among *k_pi_* and all four parameters. In particular, average habitat temperature and altitude showed a significant correlation with *k_pi_* (p < 0.01).

**Figure 2.**
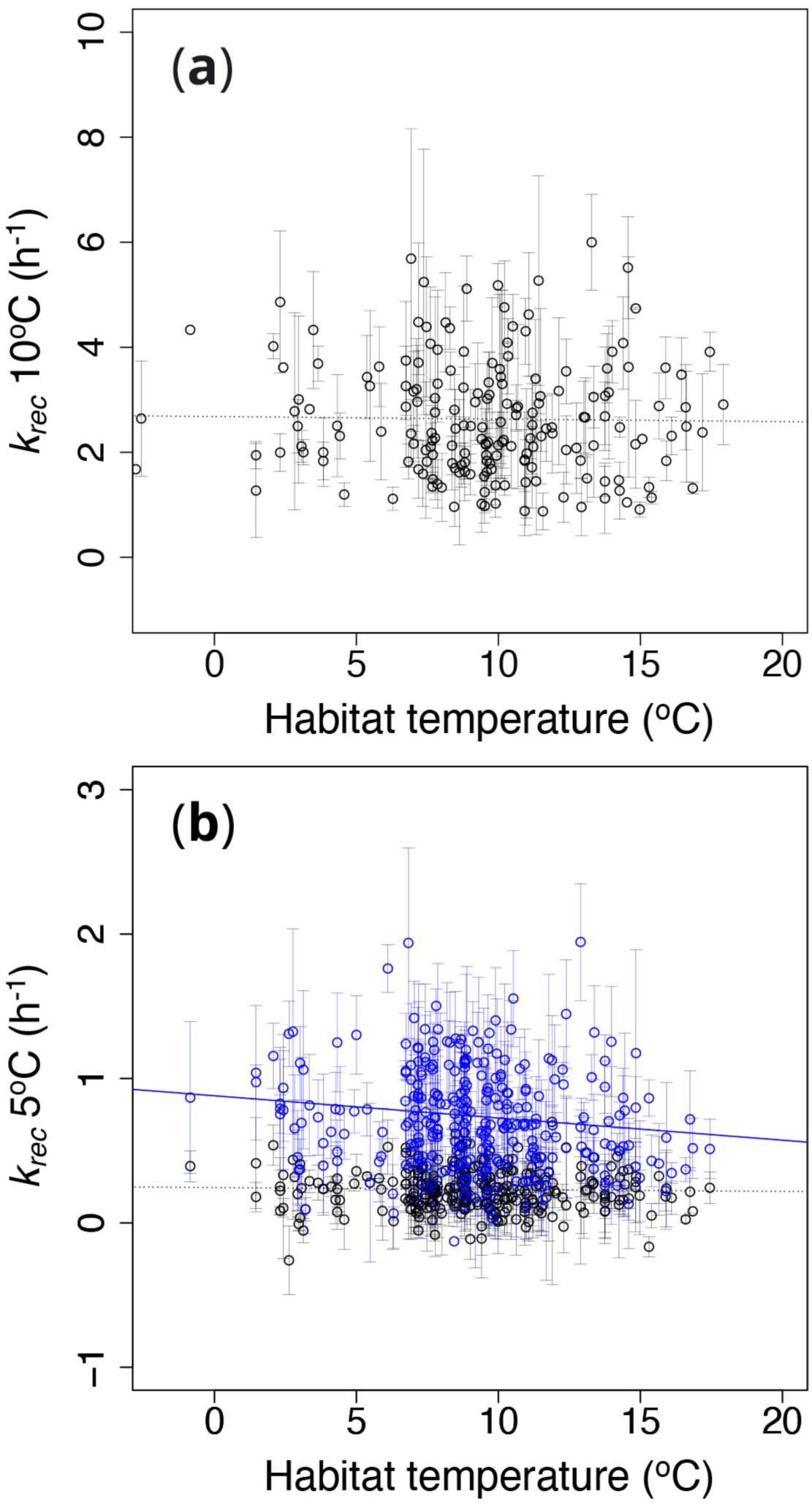
The relationships between the mean annual habitat temperature of each ecotype (°C) and the rate coefficient of PSII photoinhibition repair: *k_rec_* (h^−1^) which were obtained from photoinhibition experiments at 10 °C (a) and 5 °C (b). Each point indicates one ecotype. Plants were grown in 22°C (black symbols) or acclimated to 12°C for 3 days after grown in 22°C (blue symbols). The total length of growth was 25 days both for the control plants and the acclimated plants. Solid regression line: *y* = 0.886 – 0.016*x*, *R*^2^ = 0.024, *P* = 0.008. Dotted lines indicate statistically non-significant relationships.

**Figure 3.**
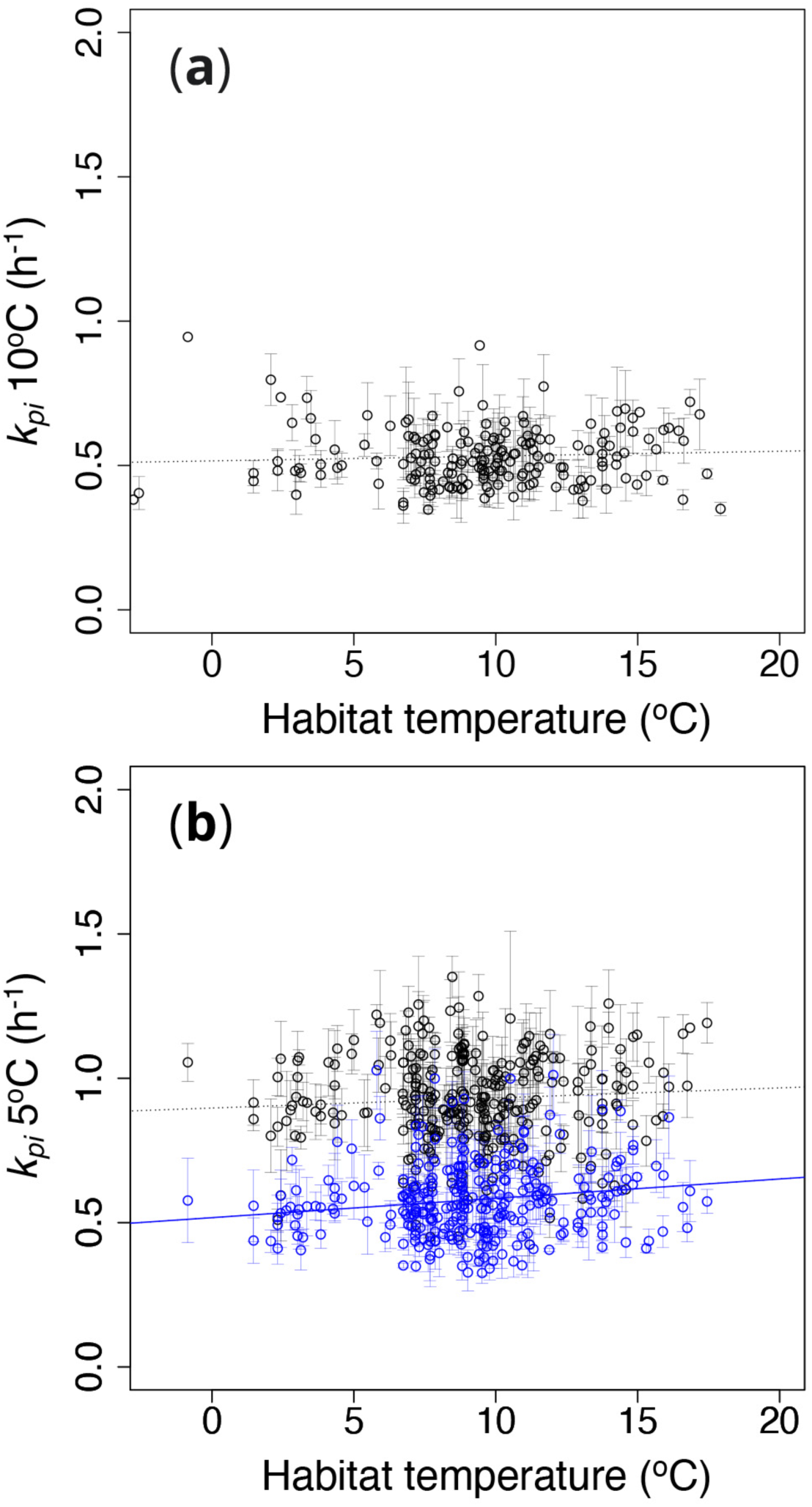
The relationships between the mean annual habitat temperature of each ecotype (°C) and the rate coefficient of PSII photoinhibition: *k_pi_* (h^−1^) which were gained from photoinhibition experiments at 10 °C (a) and 5 °C (b). Each point indicates one ecotype. Plants were grown in 22°C (black symbols) or acclimated to 12°C for 3 days after grown in 22°C (blue symbols). The total length of growth was 25 days both for the control plants and the acclimated plants. Solid regression line: *y* = 0.517 + 0.007*x*, *R*^2^ = 0.028, *P* = 0.004. Dotted lines indicate statistically non-significant relationships.

**Table 1.**
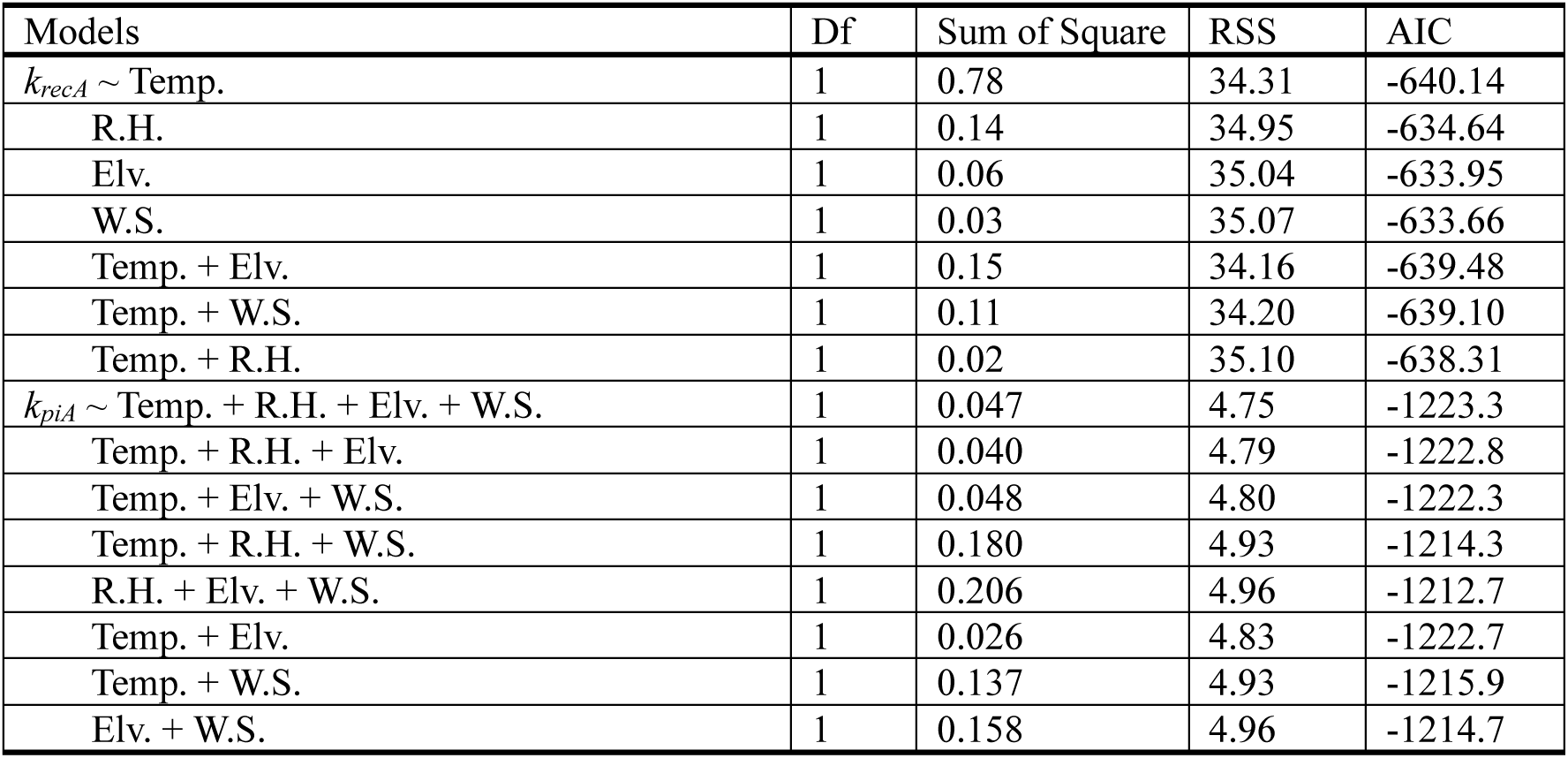
The results of the tested models of the relationships among *k_recA_* or *k_piA_* and environmental parameters (e.g. average temperature: Temp, average relative humidity: R.H., elevation: Elv. and average wind speed: W.S. of habitats) using the stepwise model selection of MASS package of R. In the case of *k_recA_*, the selected model was the relationship between *k_recA_* and average temperature (*P* = 0.00992). In the case of *k_piA_*, the selected model was the relationship between *k_piA_* and all four parameters (Temp.: *P* = 0.00043, Elv.: *P* = 0.00101, R.H.: *P* = 0.00875, W.S.: *P* = 0.11791).

There was a correlation between the rate coefficient of photoinhibition (*k_piC_*) and the rate coefficient of photoinhibition repair (*k_recC_*) at 10°C in non-hardened control plants (Figure 4a), but the correlation was not observed at 5°C both in control plants and cold-acclimated plants (Figure 4b). *k_recC_* and *k_piC_* each showed a correlation between the rate at 5°C and the rate at 10°C in non-hardened plants (Figure 4c and d). Although the cold-acclimation changed *k_rec_* and *k_pi_* remarkably as shown in Figures 2 and 3, the rate coefficient of PSII photoinhibition repair at 5°C showed a correlation between the control plants (*k_recC_*) and cold-acclimated (*k_recA_*) plants (Figure S2a). This correlation was also observed in the rate coefficients of PSII photoinhibition of control (*k_piC_*) and hardened (*k_piA_*) plants (Figure S2b). *k_recA_* at 5°C showed a correlation with *k_recC_* at 10°C (Figure S2c). The rate coefficient of PSII photoinhibition also showed this correlation (Figure S2d). There was a significant negative correlation between the indicator of the extent of non-photochemical quenching (*F_v_*/*F_m_* – (*F_v_*/*F_m_*)*_NR_*) and *k_pi_* at 5°C both in control plants and cold-acclimated plants (Figure 5a). However, (*F_v_*/*F_m_* – (*F_v_*/*F_m_*)*_NR_*) at 5°C did not show any correlation with the habitat temperature of each ecotype, even in cold-acclimated plants (Figure 5b). It is also noteworthy that *k_pi_* of control plants was higher than that of cold-acclimated plants even when they showed similar indicators of non-photochemical quenching (Figure 5a). In the case of the indicator of energy dependent quenching (*qE* indicator), it did not show any correlation with *k_pi_* nor the habitat temperature of each ecotype (Figure 5c and d).

**Figure 4.**
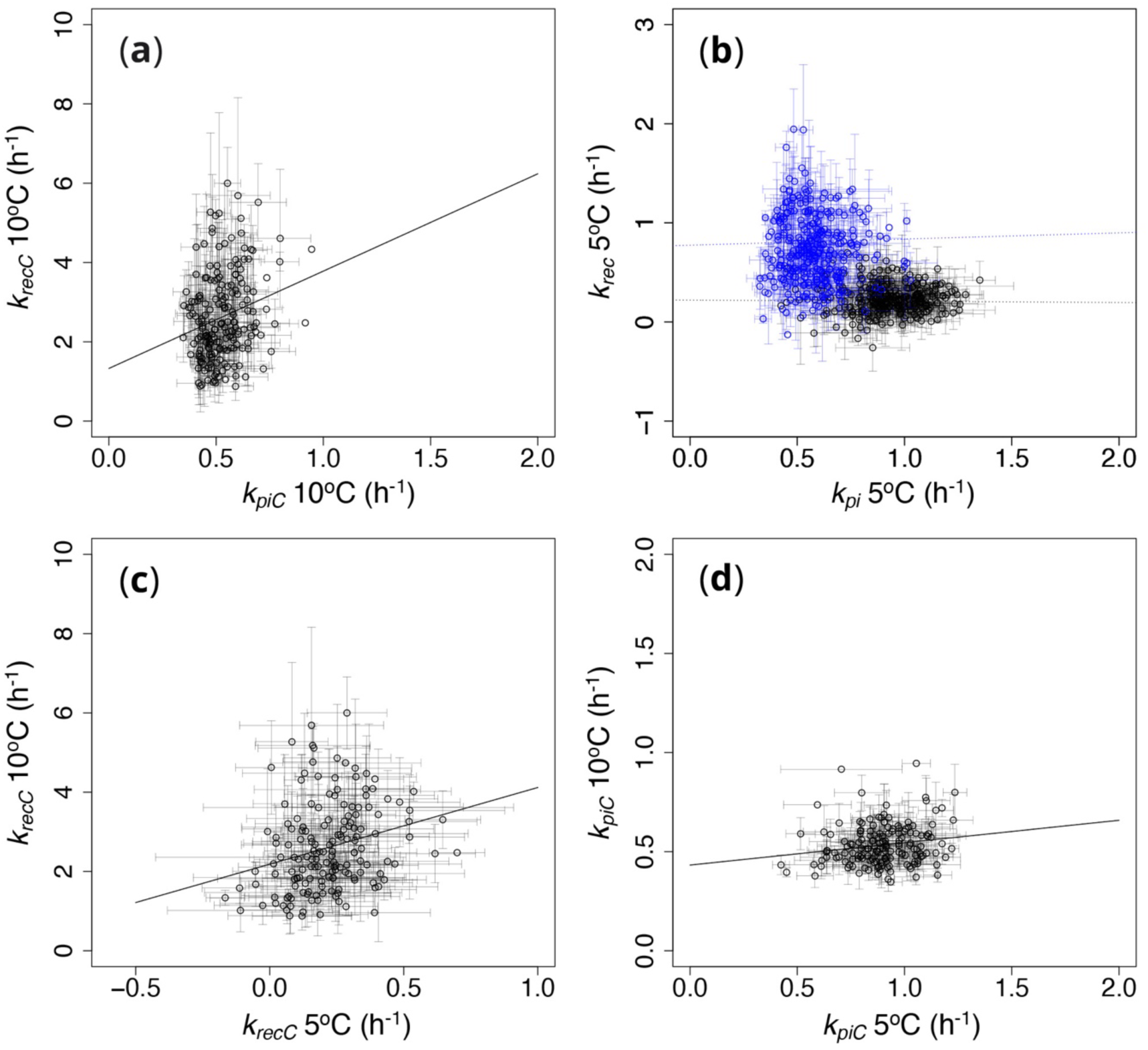
The relationships between the rate coefficient of PSII photoinhibition: *k_pi_* (h^−1^) and the rate coefficient of PSII photoinhibition repair: *k_rec_* (h^−1^) which were obtained from photoinhibition experiments at 10 °C (a) and 5 °C (b). In (b), plants were grown in 22°C (black symbols) or acclimated to 12°C for 3 days after grown in 22°C (blue symbols). The total length of growth was 25 days both for the control plants and the acclimated plants. The relationship of the rate constants of PSII photoinhibition repair: *k_rec_* (h^−1^) between at 5 °C and 10 °C (c). The relationship of the rate constants of PSII photoinhibition: *k_pi_* (h^−1^) between at 5 °C and 10 °C (d). Each point indicates one ecotype. Solid regression lines: (a) *y* = 1.33 + 2.45*x*, *R*^2^ = 0.051, *P* = 0.002; (c) *y* = 2.18 + 1.94*x*, *R*^2^ = 0.064, *P* = 0.001; (d) *y* = 0.432 + 0.113*x*, *R*^2^ = 0.03, *P* = 0.027. Dotted lines indicate statistically non-significant relationships.

**Figure 5.**
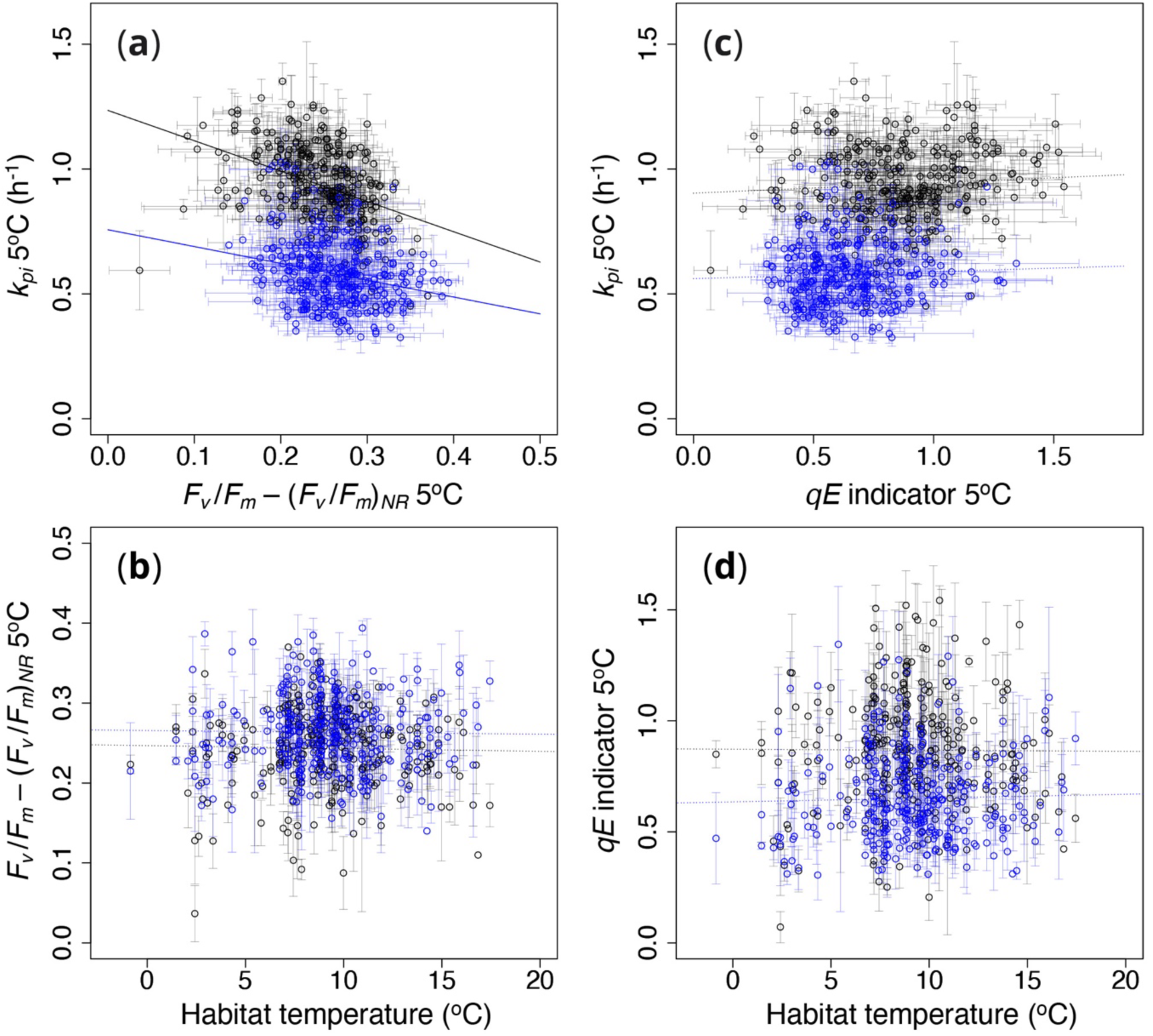
The relationships between the indicator of non-photochemical quenching (*F_v_*/*F_m_* – (*F_v_*/*F_m_*)*_NR_*) and the rate coefficient of PSII photoinhibition: *k_pi_* (h^−1^) (a). The relationships between the mean annual habitat temperature of each ecotype (°C) and the indicator of non-photochemical quenching (*F_v_*/*F_m_* – (*F_v_*/*F_m_*)*_NR_*) (b). The relationships between the indicator of energy dependent quenching (*qE* indicator) and the rate coefficient of PSII photoinhibition: *k_pi_* (h^−1^) (c). The relationships between the mean annual habitat temperature of each ecotype (°C) and the indicator of energy dependent quenching (*qE* indicator) (d). Each point indicates one ecotype. The data in this figure were obtained from photoinhibition experiments at 5 °C. Plants were grown in 22°C (black symbols) or acclimated to 12°C for 3 days after grown in 22°C (blue symbols). The total length of growth was 25 days both for the control plants and the acclimated plants. Solid regression lines: (a) control (black line) *y* = 1.22 – 0.51*x*, *R*^2^ = 0.31, *P* < 0.001; cold-acclimated (blue line) *y* = 0.857 – 0.32*x*, *R*^2^ = 0.302, *P* < 0.001. Dotted lines indicate statistically non-significant relationships.

Quantum yields of photochemical energy conversion in PSII [*Y*(*II*)] and in PSI [*Y*(*I*)] and the rate of state-transitions show dependency on treatment temperature as shown in Figure 1 b, c and d, respectively. Therefore, we also expected a correlation between the quantum yield of PSII, PSI or state-transitions rate and the habitat temperature of each ecotype. The measurement of the quantum yields and rate of state-transitions were conducted at 12°C, using 54 and 49 *A*. *thaliana* ecotypes, respectively. However, there was no statistically significant correlation in these relationships (Figure 6). Higher quantum yield of PSII and PSI may increase *k_rec_* and decrease *k_pi_* via decreasing reactive oxygen species and decreasing excess energy, respectively, but the quantum yields did not show significant correlation with *k_rec_* and *k_pi_* (Figure 7). Only the *Y*(*I*) – *Y*(*II*), which has been considered to indicate the activity of cyclic electron flow around PSI, showed a significant negative correlation with *k_rec_* (Figure 7e), but not with *k_pi_* (Figure 7f). We could not see any correlation in the relationships between the rates of state-transitions and *k_rec_* and between the rates of state-transitions and *k_pi_* (Figure 8a-d), although there was a significant correlation between the rate of state-transitions from State 1 to State 2 and that from State 2 to State 1 (Figure 8e).

**Figure 6.**
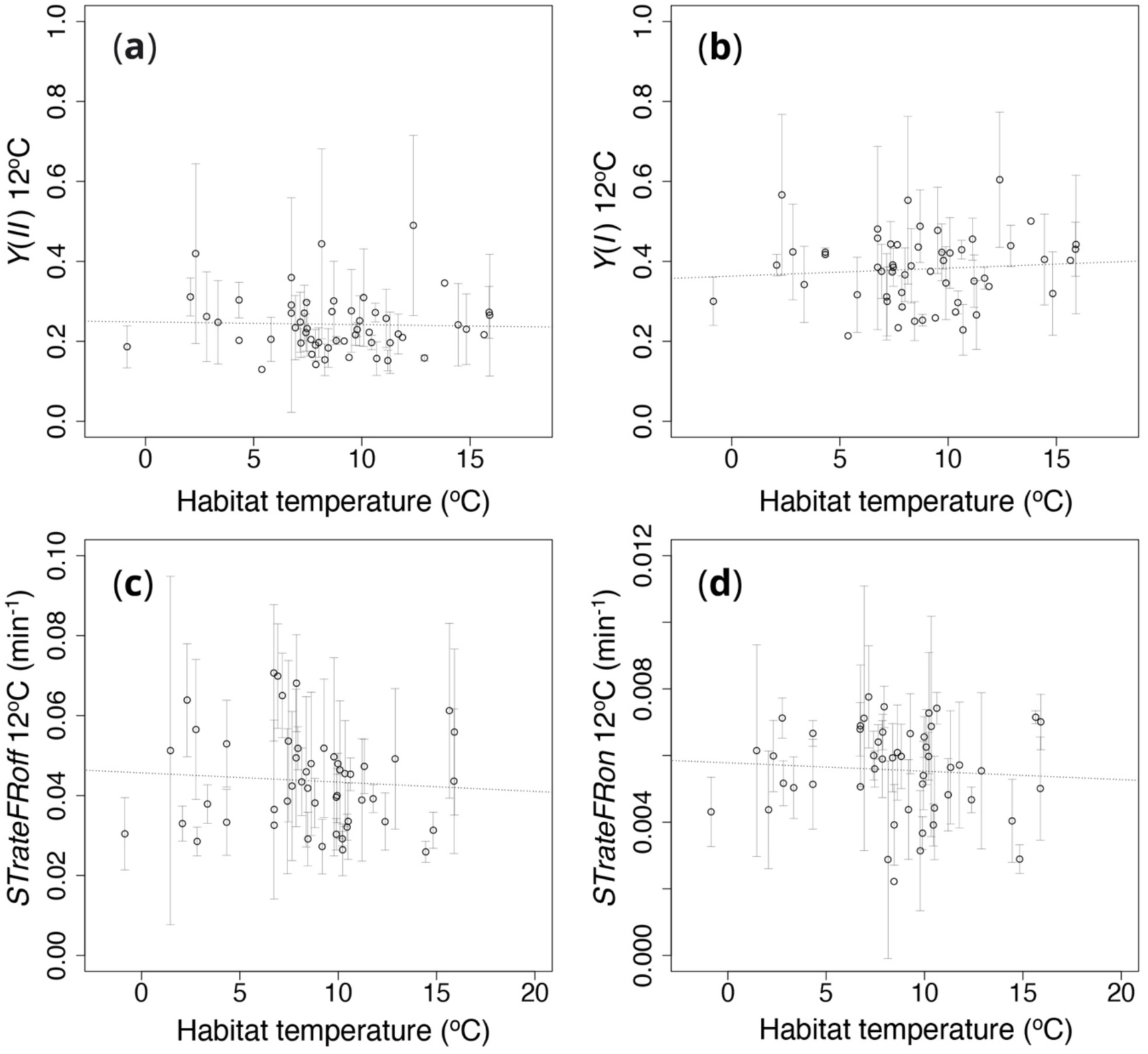
The relationships between the mean annual habitat temperature of each ecotype (°C) and (a) the quantum yield of photochemical energy conversion in PSII [*Y*(*II*)], (b) the quantum yield of photochemical energy conversion in PSI [*Y*(*I*)], (c) rate of state-transitions from State I to State II (*STrateFRoff*), and (d) rate of state-transitions from State II to State I (*STrateFRon*), which were gained at 12 °C. Each point indicates one ecotype. Dotted lines indicate statistically non-significant relationships.

**Figure 7.**
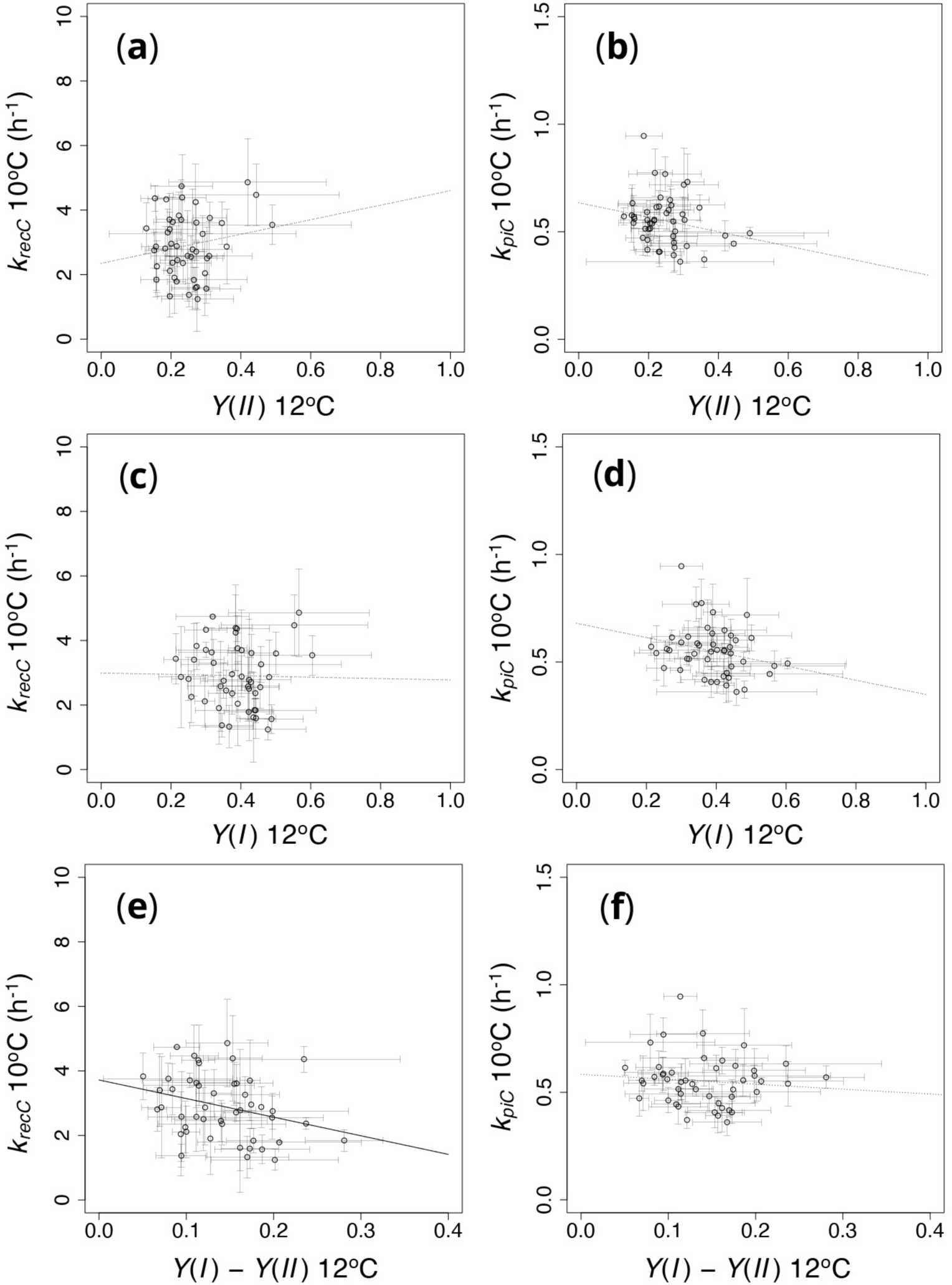
The relationships between the rate coefficient of photoinhibition repair and *Y*(*II*), *Y*(*I*) and *Y*(*I*) – *Y*(*II*) (a, c and e), and between the rate coefficient of photoinhibition and *Y*(*II*), *Y*(*I*) and *Y*(*I*) – *Y*(*II*) (b, d and f). The rate coefficients of photoinhibition and photoinhibition repair were obtained at 10°C, and *Y*(*II*) and *Y*(*I*) were obtained at 12°C. Each point indicates one ecotype. Solid regression line: (e) *y* = 3.72 – 5.76*x*, *R*^2^ = 0.087, *P* = 0.044. Dotted lines indicate statistically non-significant relationships.

**Figure 8.**
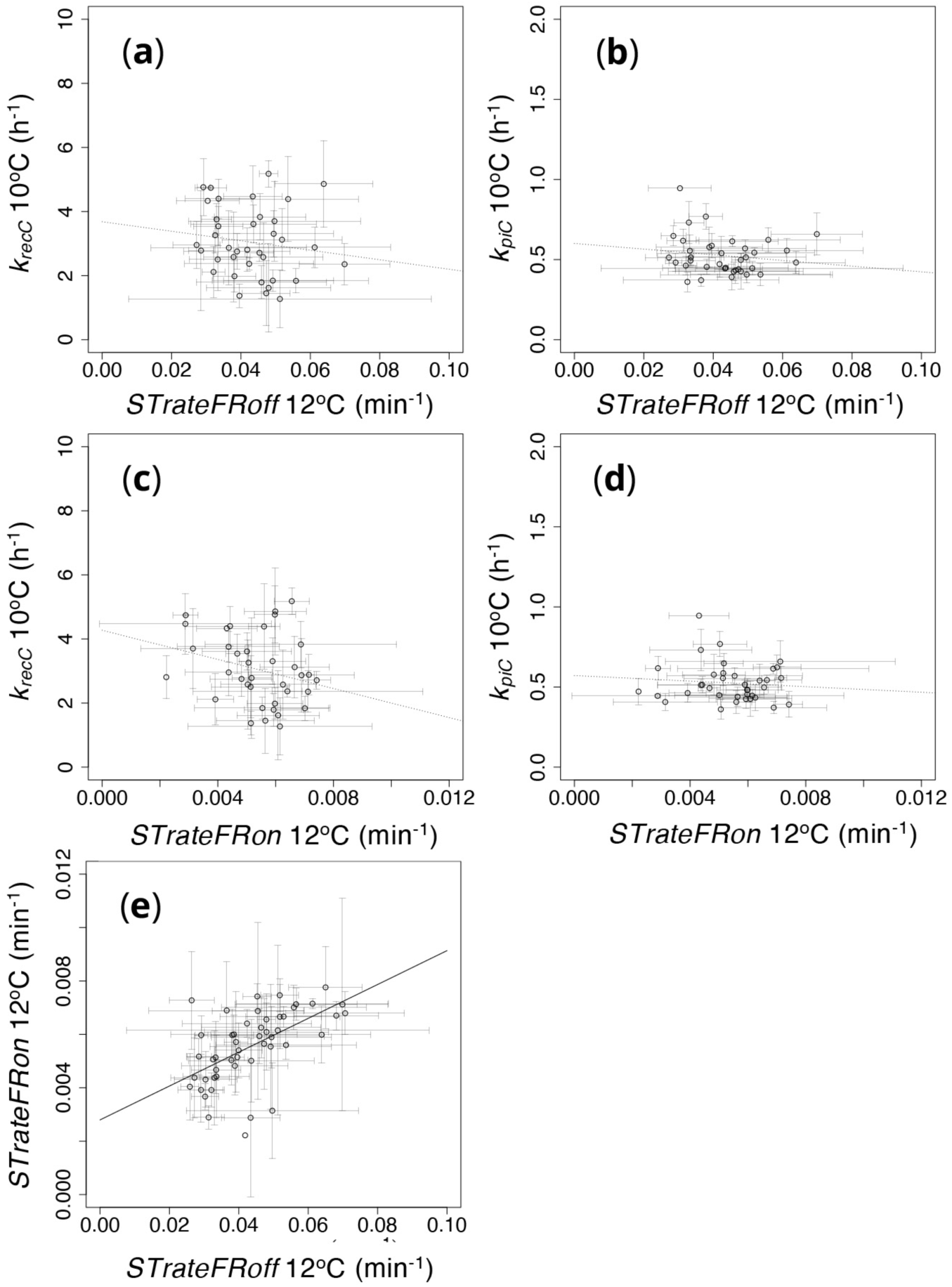
The relationships between the rate of state-transitions and the rate coefficient of photoinhibition repair (a and c) and those between the rate of state-transitions and the rate coefficient of photoinhibition (b and d). The rates of state-transitions from State I to State II (a and b) and those from State II to State I (c and d) were obtained at 12°C, and the rate coefficients of photoinhibition and photoinhibition repair were obtained at 10°C. Panel (e) shows the relationship between rate of state-transitions from State I to State II and that from State II to State I. Each point indicates one ecotype. Solid regression line: (e) *y* = 0.003 + 0.063*x*, *R*^2^ = 0.319, *P* < 0.001. Dotted lines indicate statistically non-significant relationships.

To see the effect of leaf morphological traits on the photoinhibition tolerance, we analyzed the relationship between leaf mass per area (LMA) and the rate coefficient of photoinhibition repair (*k_rec_*). *k_rec_* of control plants at 5°C showed a significant correlation with leaf mass per area: LMA (Figure 9a). The data of LMA was gained from our previous research (Oguchi *et al*., 2016), where plants were grown in ambient and elevated CO_2_ (800 ppm) conditions. *k_rec_* of control plants at 5°C also showed a significant correlation with LMA of elevated CO_2_ condition (Figure 9b). *k_rec_* of cold-acclimated plants at 5°C did not show a correlation with LMA of ambient condition, but showed a significant correlation with LMA of elevated CO_2_ condition (Figure 9c and d). It should be noted that LMA of elevated CO_2_ condition had a strong correlation with LMA of ambient condition (*R*^2^ = 0.536).

**Figure 9.**
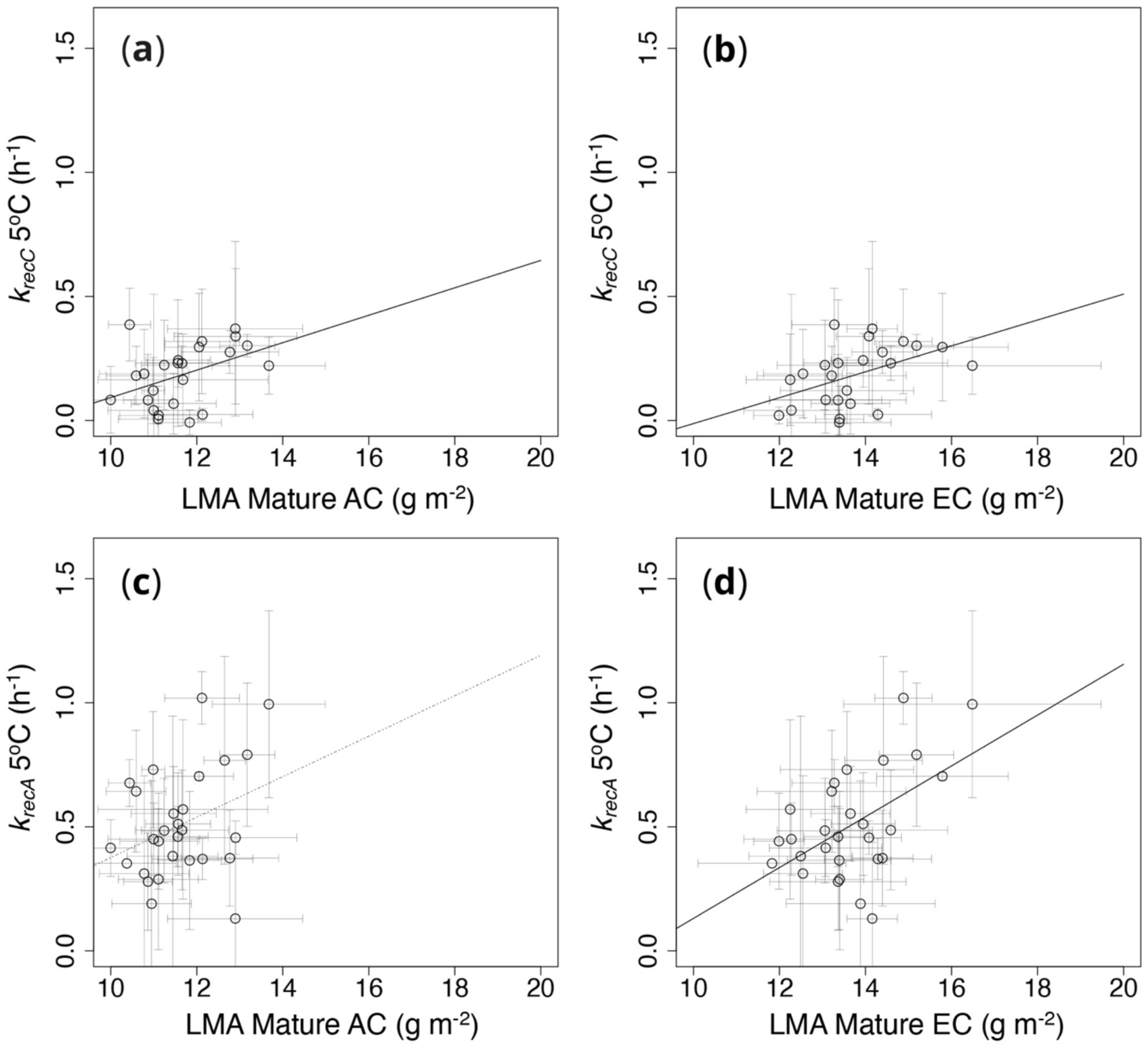
The relationships between the rate coefficient of photoinhibition repair (*k_rec_*) at 5°C and leaf mass per area (LMA). The rate coefficients of photoinhibition repair were measured with the control plants grown in 22°C (a, b) and cold-acclimated plants acclimated to 12°C for 3 days after grown in 22°C (c, d). The total length of growth was 25 days both for the control plants and the acclimated plants. We used the data of LMA of our previous report (Oguchi et al. 2016) for the analyses, where plants were grown in ambient (AC: a, c) and elevated CO_2_ (EC: b, d) conditions at 20°C for 38 days. Each point indicates one ecotype. Solid regression lines: (a) *y* = –0.461 + 0.055*x*, *R*^2^ = 0.177, *P* = 0.041; (b) *y* = –0.535 + 0.052*x*, *R*^2^ = 0.22, *P* = 0.021; (d) *y* = –0.894 + 0.102*x*, *R*^2^ = 0.271, *P* = 0.005. A dotted line indicates a statistically non-significant relationship.

## Discussion

The statistically significant correlation between the rate coefficient of photoinhibition repair of cold-acclimated plants (*k_rec_*) at 5 °C and the habitat mean temperature of each ecotype (Figure 2b) would indicate that the photoinhibition repair capacity in cold condition can affect the distribution of plants, as shown in freezing tolerance (Zhen & Ungerer, 2008; Zuther *et al*., 2012). On the other hand, it is interesting that *k_rec_* of control (non-hardened) plants did not show correlation with the habitat mean temperature, both at 5°C and 10°C (Figure 2a and b). *k_rec_* was higher in cold-acclimated plants than control plants (Figure 2b); therefore, the increase of the repair capacity in the cold-acclimation seems to be important for the fitness of *A*. *thaliana* in cold regions. Schultze & Bilger (2019) reported that Columbia ecotype of *A*. *thaliana*, grown at 21°C, increases the repair capacity of UV-B damaged photoinhibition in a low temperature (9°C) acclimation for 2-3 weeks. The result of the present study indicates that the increase in the repair capacity of photoinhibition can be stimulated by shorter (3-day) acclimation to low temperature, and that the acclimation capacity is higher in the ecotypes originating from colder habitats. The mechanism of the repair of photoinhibited PSII has been studied for long time (Gururani *et al*., 2015; Jarvi *et al*., 2015; Rantala *et al*., 2020), and it has been reported that there are multiple processes in the repair mechanism including the migration of damaged PSII complex in thylakoids, the degradation of damaged reaction center (D1 protein) and the synthesis of new reaction center, and that there are also multiple proteins facilitating the repair mechanism of PSII. However, the information about the strength of the temperature dependency of each process and the mechanism of the cold-acclimation of repair capacity is still not sufficient. It should be important to study the physiological mechanism of the enhancement of repair capacity in cold acclimation in the future.

The rate coefficient of photoinhibition of cold-acclimated plants (*k_pi_*) at 5°C also showed a significant correlation with the habitat mean temperature of each ecotype (Figure 3b), which would also support the notion that the extent of photoinhibition in cold condition can affect the distribution of plants. *k_pi_* of control plants did not show a correlation with the habitat mean temperature, as was also in the case in *k_rec_*, and *k_pi_* of cold-acclimated plants was lower than that of control plants (Figure 3a and b). Therefore, cold-acclimation would improve the fitness of *A*. *thaliana* in cold regions not only via increase in photoinhibition repair capacity but also via decrease in photoinhibition rate. It is suggested that an enhanced epidermal screening in low-temperature acclimated leaves can increase resistance to photoinhibition caused by UV-B radiation (Schultze & Bilger, 2019). The effect of screening of visible light in the cold acclimation should be also studied in the future.

A positive correlation between *k_rec_* and *k_pi_* was observed in control plants at 10°C (Figure 4a), but not observed at 5°C both in control and cold-acclimated plants (Figure 4b). According to Miyata et al. (2015), herbal species from habitats with higher light intensity showed higher *k_rec_* and lower *k_pi_*. Although, in the present study, the ecotypes from colder habitats tended to have higher *k_rec_* and lower *k_pi_* after cold-acclimation (Figures 2b and 3b), there was not a significant negative correlation between *k_rec_* and *k_pi_* even in cold-acclimated plants at 5°C, which might suggest that possessing both a high repair capacity and a low photoinhibition rate is costly. At least, it would not be simply a case of a higher repair capacity supplementing a higher rate of photoinhibition. Both *k_rec_* and *k_pi_* showed a significant correlation between the two photoinhibition temperature conditions, although *k_rec_* was markedly lower at 5°C than at 10°C (Figure 4c, d). These correlations were still observed between cold-acclimated plants and control plants (Figure S2c, d). This may be because the treatment of cold-acclimation at 12°C was relatively short (3 days) in the present research and the correlations between control plants and cold-acclimated plants were observed both in *k_rec_* and *k_pi_* (Figure S2a and b). The negative correlation between the indicator of non-photochemical quenching (*F_v_*/*F_m_* – (*F_v_*/*F_m_*)*_NR_*) and *k_pi_* both in control and cold-acclimated plants at 5°C may indicate that higher non-photochemical quenching can mitigate the rate of photoinhibition (Figure 5a). However, the lower *k_pi_* of cold-acclimated plants was not attributable to a higher non-photochemical quenching, because the difference of the indicators of non-photochemical quenching: (*F_v_*/*F_m_* – (*F_v_*/*F_m_*)*_NR_*) and *qE* indicator, were small between the control and cold-acclimated plants. Besides, both of two indicators did not show any correlation with the habitat temperature of each ecotype even in the cold-acclimated plants (Figure 5b and d). These results would indicate that, in the case of *A*. *thaliana*, the repair capacity of photoinhibition at a cold temperature is more important for survival in cold habitats than the capacity for non-photochemical quenching.

We expected negative correlations between the habitat mean temperature and quantum yield of PSII [*Y*(*II*)], quantum yield of PSI [*Y*(*I*)] and rate of state-transitions, as the supplementation for the lower activity of thylakoid membrane activities at lower temperature (Figure 1). However, none of them showed a significant correlation (Figure 6). This is similar to *k_rec_* and *k_pi_* of plants which had not been acclimated to cold-temperature (Figures 2 and 3); therefore, only cold-acclimated plants may show statistically significant correlations in these relationships, but we did not have enough time to conduct such experiments in this international collaboration because of COVID-19. We are also aware that the average annual minimum temperatures assigned to these ecotypes are presumably based on standardized atmospheric measurements that are poorly associated with soil temperature and light exposures of juvenile *A*. *thaliana* plants in situ (Geiger *et al*., 2012). The difference between the growth temperature and the habitat temperature, which is negative for ecotypes from warmer habitat and positive for ecotypes from colder habitat, may have also masked the intraspecific trend. Adams et al. (2016) showed that direction (positive or negative) of correlations between average annual temperature of ecotype origin and foliar vasculature traits differed depending on growth temperature. Therefore, it would be necessary to study the relationships between the habitat mean temperature and the photosynthetic traits studied in the present research using the ecotypes grown in different growth temperatures, in the future. Although we expected a common mechanism to maintain activities in lower temperature among the thylakoid membrane functions (i.e. photoinhibition repair, electron transport and state-transitions), there was no significant correlation among them (Figures 7 and 8) except a negative correlation between *Y*(*I*) – *Y*(*II*), which has been considered as the activity of cyclic electron flow around PSI (Kou *et al*., 2013), and *k_rec_* (Figure 7e). The activity of cyclic electron flow is considered to produce ATP even in the stressful conditions, where photosynthetic linear electron flow is low (Yamori & Shikanai, 2016). Therefore, it could help photoinhibition repair even at a low temperature (Yamori *et al*., 2011), but the observed correlation was negative. So far, we do not have enough information to explain this negative correlation. If PSII to PSI spillover is working more in the ecotypes which have higher *k_rec_* at a cold temperature, the spillover in cold condition may cause an increase in *Y*(*NA*) and, therefore, a decrease in *Y*(*I*); this may explain the lower *Y*(*I*) – *Y*(*II*) values of the ecotypes having higher *k_rec_*.

Because it is well known that leaf mass per area (LMA) shows light acclimation in response to different light intensities with higher LMA in higher light intensity (Björkman, 1981; Poorter *et al*., 2019) and that LMA is one of the important traits which shows a correlation with plant strategies as suggested in Leaf Economic Spectrum theory (Wright *et al*., 2004; Diaz *et al*., 2022), we analyzed the relationships between the rate coefficient of photoinhibition repair and LMA (Figure 9). Although we used the data of LMA gained in our previous study (Oguchi *et al*., 2016), where the growth condition differs in some aspects, we were still able to see significant correlations. We think this positive correlation between the repair capacity and LMA can be related to the previous reports which showed that LMA can be increased at lower growth temperature in *A*. *thaliana* (Gorsuch *et al*., 2010) and in many more species (Cohu *et al*., 2014; Read *et al*., 2014). It would be important to study the meaning of this correlation in the future.

## Conclusion

The rate coefficient of photoinhibition repair (*k_rec_*) can be increased by the cold-acclimation even in a few days. The ecotypes of *A*. *thaliana* from colder regions have a higher capacity for photoinhibition repair and lower rate of photoinhibition after the cold-acclimation. This should indicate that photoinhibition tolerance can affect the distribution of plants, especially in colder regions.

## Acknowledgements

The present study was supported by JSPS Grants-in-Aid for Scientific Research (KAKENHI) Grant Number 17KK0142 (Fund for the Promotion of Joint International Research) and its original project Grant Number JP16K18614 (General). Authors appreciate the help of Dr. Rhys Wyber and Dr. Zbigniew Kolber for setting up LIFT instruments. Authors also thank Dr. Mengmeng Zhang, Lukas Bender, Jenny Rath and Christine Larsen for their assistance in growing plants and conducting experiments, and Prof. Takashi Tsuchimatsu for discussing the selection of *Arabidopsis* ecotypes. We thank Prof. Sharon Robinson, University of Wollongong NSW 2522, for access to the LIFT and ASRM instruments.

## Author contributions

RO planned and designed the research; RO, SN, AP, HO, BO and WSC performed experiments; RO, SN, AP, HO, KH and BO analyzed data; and RO, BO and WSC wrote the manuscript.

## Data availability

The data that support the findings of this study are available from the corresponding author upon reasonable request.

