## Supplementary information for "A negative correlation between the rate coefficient of repair after photoinhibition of cold acclimated plants and the mean annual temperature of the habitats of *Arabidopsis thaliana* ecotypes"

The following Supplementary Information is available for this article:

**Fig. S1** An example of the method to calculate the rates of state-transition using the Light Induced Fluorescence Transient (LIFT).

**Fig. S2** The relationship of the rate coefficient of PSII photoinhibition repair: $k_{rec}$ ($h^{-1}$) (a) and the rate coefficient of PSII photoinhibition: $k_{pi}$ ($h^{-1}$) (b) at 5 °C between the control plants (grown in 22°C) and cold-acclimated plants (acclimated to 12°C for 3 days after grown in 22°C). The relationship of the rate coefficient of PSII photoinhibition repair: $k_{rec}$ ($h^{-1}$) (c) and the rate coefficient of PSII photoinhibition: $k_{pi}$ ($h^{-1}$) (d) between the control plants photoinhibited at 10°C and cold-acclimated plants photoinhibited at 5 °C.

**Table S1** List of the *Arabidopsis thaliana* ecotypes.

**Methods S1** Equations of energy-dependent non-photochemical quenching.

**Fig. S1** An example of the method to calculate the rates of state-transition using the Light Induced Fluorescence Transient (LIFT). In the case of calculating the rate of state-transition after far-red light off (*STrateFRoff*), only the points larger than {"$F_m$' before FR on" + 0.25 ・ ("Max $F_m$' after FR off" - "$F_m$' before FR on")} were used to avoid the tailing effect on the slope.

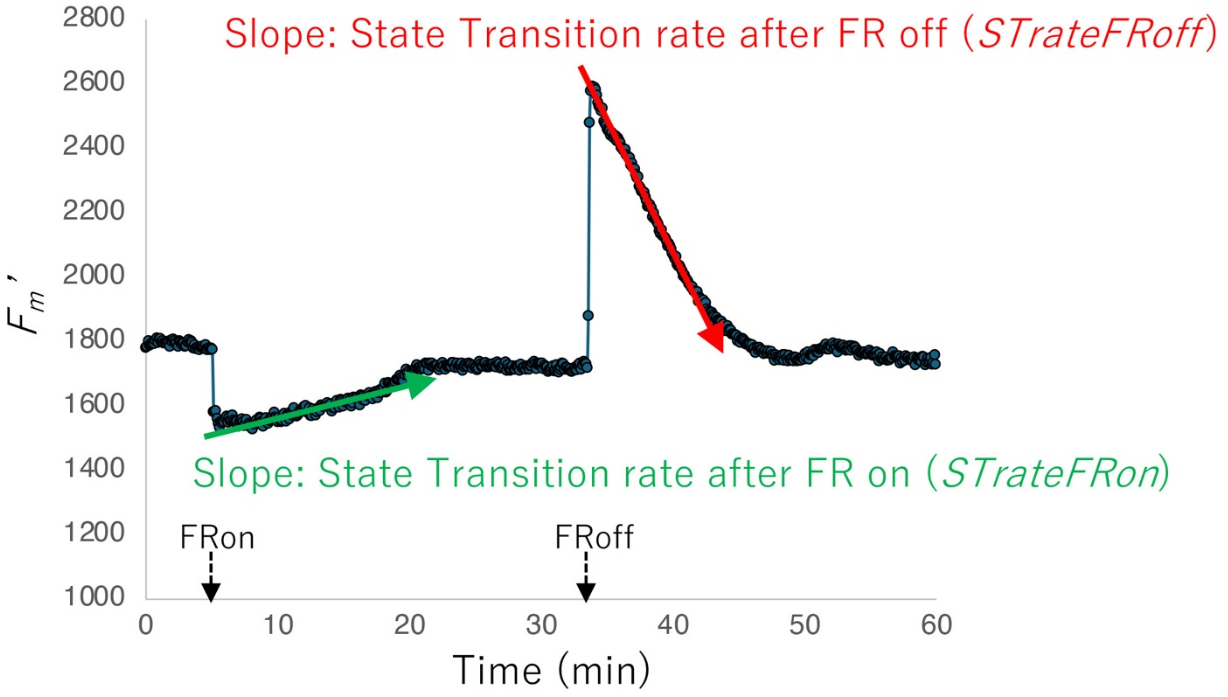

**Fig. S2** The relationship of the rate coefficient of PSII photoinhibition repair: $k_{rec}$ ($h^{-1}$) (a) and the rate coefficient of PSII photoinhibition: $k_{pi}$ ($h^{-1}$) (b) at 5 °C between the control plants (grown in 22°C) and cold-acclimated plants (acclimated to 12°C for 3 days after grown in 22°C). The relationship of the rate coefficient of PSII photoinhibition repair: $k_{rec}$ ($h^{-1}$) (c) and the rate coefficient of PSII photoinhibition: $k_{pi}$ ($h^{-1}$) (d) between the control plants photoinhibited at 10°C and cold-acclimated plants photoinhibited at 5 °C. Solid regression lines: (a) $y = 0.566 + 0.743x$, $R^2 = 0.097$, $P < 0.001$; (b) $y = 0.214 + 0.392x$, $R^2 = 0.23$, $P < 0.001$; (c) $y = 0.51 + 0.043x$, $R^2 = 0.03$, $P = 0.027$; (d) $y = 0.363 + 0.312x$, $R^2 = 0.144$, $P < 0.001$.

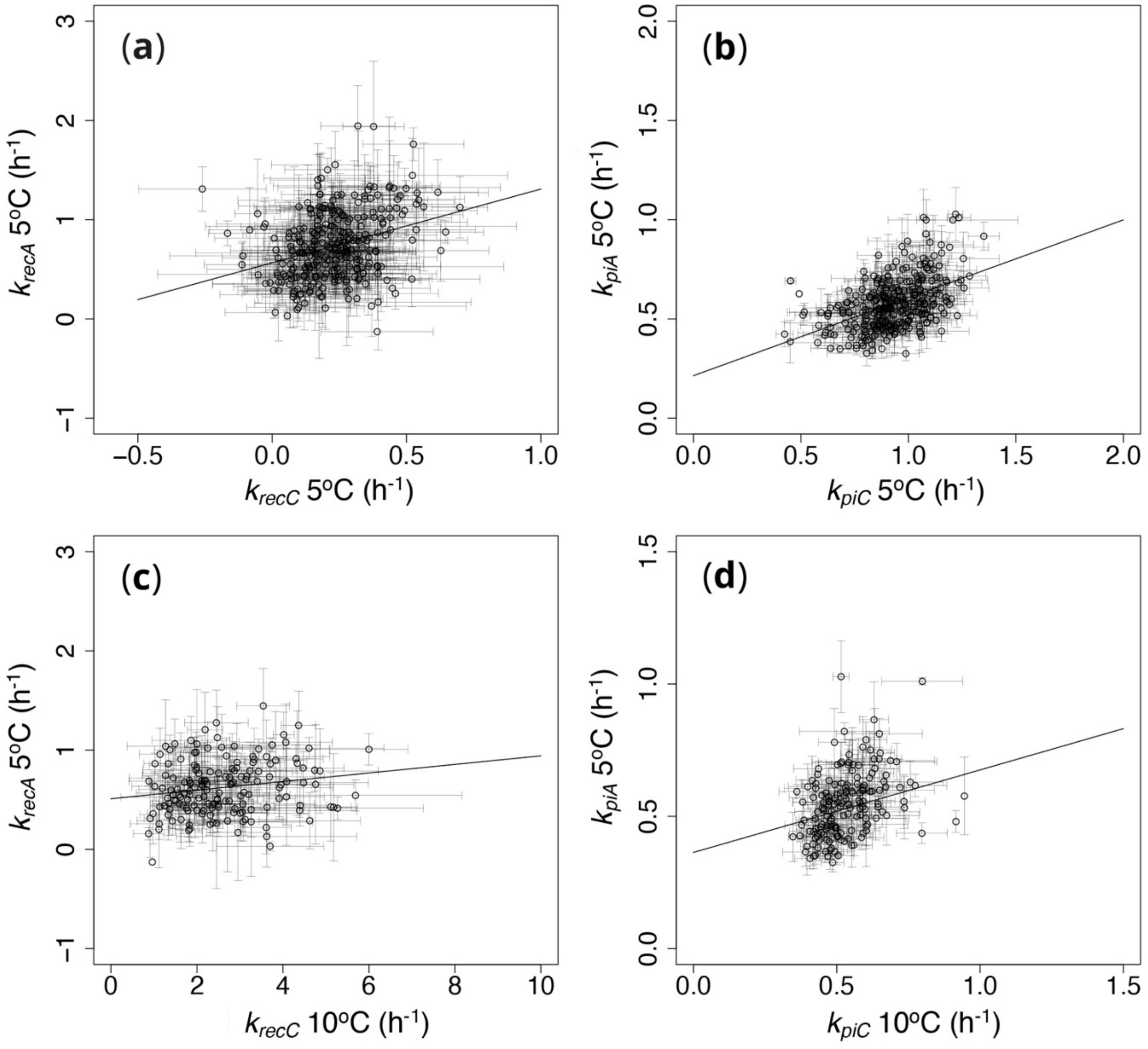

**Table S1** List of the *Arabidopsis thaliana* ecotypes. The ecotypes used for photoinhibition measurements (PI 5ºC and PI 10ºC), photosystems-activity measurement (PS act.) and state-transitions measurement (ST) are shown by circles. West longitudes are shown as negative value.

| ABRC number | List name | Country | Latitude | Longitude | PI 5ºC | PI 10ºC | PS act. | ST |
|---|---|---|---|---|---|---|---|---|
| CS76347 | Aitba-2 | Morocco | 31.5 | -7.5 | ○ | | ○ | ○ |
| CS76476 | Dra-0 | Czech | 49.4 | 16.3 | ○ | ○ | ○ | ○ |
| CS76486 | Et-0 | France | 44.6 | 2.6 | ○ | ○ | ○ | ○ |
| CS76534 | Krot-0 | Germany | 49.6 | 11.6 | ○ | ○ | ○ | ○ |
| CS76560 | Neo-6 | Tajikistan | 37.4 | 72.5 | ○ | ○ | ○ | ○ |
| CS76667 | App1-12 | Sweden | 56.3 | 16.0 | ○ | | ○ | ○ |
| CS76693 | Basta-3 | Russia | 51.8 | 79.5 | ○ | ○ | ○ | ○ |
| CS76712 | Bisig-1 | Italy | 39.5 | 16.3 | ○ | ○ | ○ | ○ |
| CS76721 | IP-Bra-0 | Spain | 42.5 | -6.2 | ○ | ○ | | ○ |
| CS76774 | IP-Cmo-3 | Spain | 40.1 | -4.7 | ○ | ○ | ○ | ○ |
| CS76804 | Dolna-1-39 | Bulgaria | 42.3 | 23.1 | ○ | ○ | ○ | ○ |
| CS76806 | Dor-10 | Sweden | 63.0 | 17.5 | ○ | ○ | ○ | ○ |
| CS76887 | Gradi-1 | Croatia | 45.2 | 18.7 | ○ | ○ | | ○ |
| CS76891 | Gron 12 | Sweden | 62.8 | 18.2 | ○ | ○ | ○ | ○ |
| CS76934 | Hov3-2 | Sweden | 56.1 | 13.7 | ○ | ○ | ○ | ○ |
| CS76939 | IP-Hoy-0 | Spain | 40.4 | -5.0 | ○ | ○ | ○ | ○ |
| CS76964 | Kavlinge-1 | Sweden | 55.8 | 13.1 | ○ | ○ | ○ | ○ |
| CS76979 | Kolyv-5 | Russia | 51.3 | 82.6 | ○ | ○ | | ○ |
| CS76984 | Krazo-1 | Russia | 53.1 | 52.0 | ○ | ○ | ○ | ○ |
| CS76992 | KYC-33 | USA | 37.9 | -84.5 | ○ | ○ | ○ | ○ |
| CS77015 | Lebja-1 | Russia | 51.7 | 80.8 | ○ | | | ○ |
| CS77021 | Ler-1 | Germany | 48.0 | 10.9 | ○ | | ○ | ○ |
| CS77026 | Lerik2-3 | Azerbaijan | 38.8 | 48.6 | ○ | ○ | ○ | ○ |
| CS77034 | Lesno-4 | Russia | 53.0 | 52.0 | ○ | | ○ | ○ |
| CS77044 | Lis-3 | Sweden | 56.0 | 14.8 | ○ | | ○ | ○ |
| CS77071 | Marce-1 | Italy | 38.9 | 16.5 | ○ | ○ | ○ | ○ |
| CS77104 | IP-Moe-0 | Spain | 41.8 | 2.4 | ○ | | | ○ |
| CS77176 | IP-Pie-0 | Spain | 40.5 | -5.3 | ○ | | | ○ |
| CS77229 | IP-Sac-0 | Spain | 42.1 | -6.7 | ○ | ○ | | ○ |
| CS77364 | TFa 08 | Sweden | 63.0 | 18.3 | ○ | ○ | ○ | ○ |

Table S1 continued

| ABRC number | List name | Country | Latitude | Longitude | PI 5°C | PI 10°C | PS act. | ST |
|---|---|---|---|---|---|---|---|---|
| CS77383 | TOU-A1-89 | France | 46.7 | 4.1 | ○ | ○ | | ○ |
| CS78772 | TV-7 | Sweden | 55.6 | 14.3 | ○ | ○ | ○ | ○ |
| CS78777 | Uk-3 | Germany | 48.0 | 7.8 | ○ | | | ○ |
| CS78780 | UKID116 | UK | 56.7 | -6.0 | ○ | | | ○ |
| CS78781 | Cal-2 | UK | 53.3 | -1.6 | ○ | ○ | ○ | ○ |
| CS78789 | UKID74 | UK | 51.0 | -3.1 | ○ | ○ | ○ | ○ |
| CS78805 | UKSE06-500 | UK | 51.1 | 0.6 | ○ | | | ○ |
| CS78809 | UKSW06-207 | UK | 50.4 | -4.9 | ○ | | | ○ |
| CS78811 | UKSW06-285 | UK | 50.3 | -4.9 | ○ | ○ | ○ | ○ |
| CS78834 | Vastervik | Sweden | 57.8 | 16.6 | ○ | | | ○ |
| CS78835 | IP-Vav-0 | Portugal | 38.5 | -8.0 | ○ | ○ | ○ | ○ |
| CS78836 | IP-Vaz-0 | Spain | 42.3 | -3.0 | ○ | ○ | ○ | ○ |
| CS78846 | IP-Vin-0 | Spain | 42.8 | -5.8 | ○ | ○ | | ○ |
| CS78854 | WAV-8 | France | 50.7 | 3.0 | ○ | ○ | | ○ |
| CS78876 | ZdrI 2-21 | Czech | 49.4 | 16.3 | ○ | ○ | ○ | ○ |
| CS78891 | Castelfed-4-210 | Italy | 46.3 | 11.3 | ○ | ○ | | ○ |
| CS78901 | Gy-0 | France | 49.0 | 2.0 | ○ | ○ | ○ | ○ |
| CS78956 | LI-SET-019 | USA | 40.9 | -73.1 | ○ | ○ | ○ | ○ |
| CS78969 | KBS-Mac-74 | USA | 42.4 | -85.4 | ○ | ○ | ○ | ○ |
| CS79015 | HS-12 | USA | 42.4 | -71.1 | ○ | ○ | ○ | ○ |
| CS1084 | Co-1 | Portugal | 40.5 | -8.5 | ○ | ○ | | |
| CS1286 | Kn-0 | Lithuania | 54.5 | 23.5 | ○ | | | |
| CS1306 | Lc-0 | UK | 57.0 | -4.0 | ○ | ○ | | |
| CS1566 | Tu-0 | Italy | 45.0 | 7.0 | ○ | ○ | | |
| CS1594 | Wil-1 | Russia | 55.0 | 25.0 | ○ | ○ | | |
| CS6855 | Sf-1 | Spain | 41.0 | 3.0 | ○ | ○ | | |
| CS8069 | Santa Clara | USA | 37.0 | -121.0 | ○ | ○ | | |
| CS8143 | WAR | USA | 41.0 | -71.0 | ○ | ○ | | |
| CS22456 | Sapporo-0 | Japan | 43.0 | 141.0 | ○ | ○ | | |
| CS22525 | Tiv-1 | Italy | 42.0 | 12.7 | ○ | ○ | | |

Table S1 continued

| ABRC number | List name | Country | Latitude | Longitude | PI 5ºC | PI 10ºC | PS act. | ST |
|---|---|---|---|---|---|---|---|---|
| CS22623 | Ws-0 | Russia | 52.0 | 30.0 | ○ | ○ | | |
| CS22651 | Kondara | Tajikistan | 38.0 | 68.0 | ○ | ○ | | |
| CS22661 | Nz1 | New Zealand | 37.0 | 175.0 | ○ | ○ | | |
| CS22671 | Aur-0 | USA | 45.0 | -122.0 | ○ | ○ | | |
| CS22676 | Bay-0 | Germany | 49.5 | 11.0 | ○ | ○ | | |
| CS22679 | Bur-0 | Ireland | 53.5 | -6.0 | ○ | ○ | | |
| CS22681 | Col-0 | USA | 38.0 | -92.0 | ○ | ○ | | |
| CS22686 | Ler-0 | Germany | 48.0 | 8.0 | ○ | ○ | | |
| CS22688 | Rrs-7 | USA | 41.0 | -86.0 | ○ | ○ | | |
| CS22690 | Sha | Tajikistan | 38.0 | 68.0 | ○ | ○ | | |
| CS22691 | Tamm-2 | Finland | 59.0 | 23.0 | ○ | ○ | | |
| CS22693 | Tsu-1 | Japan | 34.0 | 136.0 | ○ | ○ | | |
| CS22765 | Yeg-1 | Armenia | 39.9 | 45.4 | ○ | | | |
| CS28195 | Ct-1 | Italy | 37.0 | 15.0 | ○ | ○ | | |
| CS75700 | Olympia-1 | Greece | 37.0 | 21.0 | ○ | ○ | | |
| CS75715 | Schie-1 | Germany | 51.5 | 10.4 | ○ | ○ | | |
| CS75719 | Tanz-1 | Tanzania | 2.9 | 36.0 | ○ | ○ | | |
| CS76355 | Castelfed-4-212 | Italy | 46.3 | 11.3 | ○ | | | |
| CS76367 | Lago-1 | Italy | 39.2 | 16.3 | ○ | | | |
| CS76368 | Apost-1 | Italy | 39.0 | 16.5 | ○ | | | |
| CS76372 | Jablo-1 | Bulgaria | 41.6 | 25.2 | ○ | ○ | | |
| CS76377 | Stepn-2 | Russia | 54.1 | 60.5 | ○ | ○ | | |
| CS76380 | Sij-2 | Uzbekistan | 41.5 | 70.1 | ○ | ○ | | |
| CS76387 | Xan-1 | Azerbaijan | 38.7 | 48.8 | ○ | ○ | | |
| CS76395 | Kastel-1 | Ukraine | 44.6 | 34.4 | ○ | | | |
| CS76406 | Rue3-1-31 | Germany | 48.6 | 9.2 | ○ | | | |
| CS76407 | TueV-13 | Germany | 48.5 | 9.1 | ○ | | | |
| CS76409 | Agu-1 | Spain | 41.3 | -1.3 | ○ | ○ | ○ | |
| CS76417 | Qui-0 | Spain | 42.7 | -6.9 | ○ | | | |
| CS76420 | Copac-1 | Romania | 46.1 | 22.0 | ○ | ○ | | |

Table S1 continued

| ABRC number | List name | Country | Latitude | Longitude | PI 5ºC | PI 10ºC | PS act. | ST |
|---|---|---|---|---|---|---|---|---|
| CS76432 | Alst-1 | UK | 54.8 | -2.4 | ○ | ○ | | |
| CS76444 | Bch-1 | Germany | 49.5 | 9.3 | ○ | ○ | ○ | |
| CS76450 | Bl-1 | Italy | 44.5 | 11.3 | ○ | ○ | | |
| CS76464 | Chi-0 | Russia | 53.8 | 34.7 | ○ | | | |
| CS76468 | Co-1 | Portugal | 40.1 | -8.3 | ○ | ○ | | |
| CS76469 | Com-1 | France | 49.4 | 2.8 | ○ | | | |
| CS76485 | Est | Estonia | 58.7 | 25.0 | ○ | ○ | ○ | |
| CS76490 | Ga-0 | Germany | 50.3 | 8.0 | ○ | | | |
| CS76493 | Gie-0 | Germany | 50.6 | 8.7 | ○ | | | |
| CS76498 | Gu-0 | Germany | 50.3 | 8.0 | ○ | | | |
| CS76519 | Jl-3 | Czech | 49.2 | 16.6 | ○ | | | |
| CS76526 | Kil-0 | UK | 55.6 | -5.7 | ○ | | | |
| CS76532 | Kondara | Tajikistan | 38.5 | 68.5 | ○ | | | |
| CS76535 | Kyoto | Japan | 35.0 | 135.8 | ○ | | | |
| CS76538 | La-0 | Poland | 52.7 | 15.2 | ○ | ○ | | |
| CS76545 | Lm-2 | France | 48.0 | 0.5 | ○ | ○ | ○ | |
| CS76555 | Ms-0 | Russia | 55.8 | 37.6 | ○ | | | |
| CS76557 | Mz-0 | Germany | 50.3 | 8.3 | ○ | | | |
| CS76562 | Nok-3 | Netherlands | 52.2 | 4.5 | ○ | | | |
| CS76576 | Pog-0 | Canada | 49.3 | -123.2 | ○ | ○ | | |
| CS76579 | Pu2-23 | Czech | 49.4 | 16.4 | ○ | | | |
| CS76582 | Ra-0 | France | 46.0 | 3.3 | ○ | | | |
| CS76592 | RRS-10 | USA | 41.6 | -86.4 | ○ | | | |
| CS76599 | Sei-0 | Italy | 46.5 | 11.6 | ○ | ○ | ○ | |
| CS76608 | Ta-0 | Czech | 49.5 | 14.5 | ○ | | | |
| CS76609 | Tac-0 | USA | 47.2 | -122.5 | ○ | | | |
| CS76611 | Tha-1 | Netherlands | 52.1 | 4.3 | ○ | ○ | | |
| CS76614 | Tol-0 | USA | 41.7 | -83.6 | ○ | | | |
| CS76623 | Van-0 | Canada | 49.3 | -123.2 | ○ | ○ | | |
| CS76628 | Wei-0 | Switzerland | 47.3 | 8.3 | ○ | ○ | | |

Table S1 continued

| ABRC number | List name | Country | Latitude | Longitude | PI 5ºC | PI 10ºC | PS act. | ST |
|---|---|---|---|---|---|---|---|---|
| CS76645 | Adam-1 | Russia | 51.4 | 60.0 | ○ | ○ | | |
| CS76652 | Ale-Stenar-44-4 | Sweden | 55.4 | 14.1 | ○ | ○ | | |
| CS76653 | Ale-Stenar-56-14 | Sweden | 55.4 | 14.1 | ○ | | ○ | |
| CS76668 | App1-14 | Sweden | 56.3 | 16.0 | ○ | | | |
| CS76669 | App1-16 | Sweden | 56.3 | 16.0 | ○ | | | |
| CS76670 | IP-Ara-4 | Spain | 41.7 | -3.7 | ○ | | | |
| CS76671 | IP-Are-0 | Spain | 41.0 | -4.7 | ○ | | | |
| CS76674 | IP-Aru-0 | Spain | 41.8 | 2.5 | ○ | | | |
| CS76679 | Bach-7 | Germany | 48.4 | 8.8 | ○ | | | |
| CS76685 | Bak-5 | Georgia | 41.8 | 43.5 | ○ | | | |
| CS76689 | IP-Bar-1 | Spain | 41.4 | 2.1 | ○ | ○ | | |
| CS76690 | Basen-1 | Italy | 40.4 | 16.8 | ○ | ○ | | |
| CS76696 | Bela-1 | Slovakia | 48.5 | 18.9 | ○ | | | |
| CS76700 | IP-Ben-0 | Spain | 38.4 | -2.7 | ○ | | | |
| CS76701 | Berg-1 | Germany | 48.4 | 8.8 | ○ | | | |
| CS76729 | Brosarp-21-140 | Sweden | 55.7 | 14.1 | ○ | ○ | | |
| CS76734 | Bur-0 | Ireland | 53.1 | -9.1 | ○ | | | |
| CS76739 | IP-Cad-0 | Spain | 40.4 | -5.7 | ○ | | | |
| CS76743 | IP-Cas-0 | Spain | 38.5 | -3.4 | ○ | ○ | | |
| CS76744 | Castelfed-1-195 | Italy | 46.3 | 11.3 | ○ | | | |
| CS76745 | Castelfed-1-196 | Italy | 46.3 | 11.3 | ○ | ○ | | |
| CS76748 | Castelfed-1-199 | Italy | 46.3 | 11.3 | ○ | | | |
| CS76754 | Castelfed-3-205 | Italy | 46.3 | 11.3 | ○ | ○ | | |
| CS76756 | Castelfed-3-207 | Italy | 46.3 | 11.3 | ○ | | | |
| CS76757 | Castelfed-3-208 | Italy | 46.3 | 11.3 | ○ | | | |
| CS76761 | IP-Cdc-3 | Spain | 41.2 | -4.5 | ○ | | | |
| CS76769 | Choto-1 | Bulgaria | 41.5 | 23.3 | ○ | ○ | | |
| CS76777 | IP-Cod-0 | Spain | 41.3 | -1.3 | ○ | ○ | | |
| CS76779 | CSHL-5 | USA | 40.9 | -73.5 | ○ | ○ | | |
| CS76782 | IP-Cor-0 | Spain | 40.8 | -2.0 | ○ | | | |

Table S1 continued

| ABRC number | List name | Country | Latitude | Longitude | PI 5°C | PI 10°C | PS act. | ST |
|---|---|---|---|---|---|---|---|---|
| CS76786 | Ct-1 | Italy | 37.3 | 15.0 | ○ | | | |
| CS76791 | Da-0 | Germany | 49.9 | 8.7 | ○ | ○ | | |
| CS76808 | Doubravnik7 | Czech | 49.4 | 16.3 | ○ | | | |
| CS76812 | Draha2 | Czech | 49.4 | 16.3 | ○ | | | |
| CS76815 | DraIII-1 | Czech | 49.4 | 16.3 | ○ | | | |
| CS76816 | DraIV 1-11 | Czech | 49.4 | 16.3 | ○ | ○ | ○ | |
| CS76826 | Eden-1 | Sweden | 62.9 | 18.2 | ○ | | | |
| CS76844 | Epidauros-1 | Greece | 37.6 | 23.1 | ○ | ○ | | |
| CS76846 | IP-Esn-2 | Spain | 42.3 | 0.2 | ○ | ○ | ○ | |
| CS76852 | FaL 1 | Sweden | 63.0 | 18.3 | ○ | | | |
| CS76885 | Gr-5 | Austria | 47.0 | 15.5 | ○ | | | |
| CS76890 | Groch-1 | Bulgaria | 41.7 | 24.4 | ○ | ○ | | |
| CS76897 | H55 | Czech | 49.0 | 15.0 | ○ | ○ | ○ | |
| CS76902 | Ha-P2-1 | Germany | 48.5 | 9.0 | ○ | | | |
| CS76906 | Hadd-3 | Sweden | 57.3 | 15.9 | ○ | ○ | | |
| CS76909 | Halca-1 | Slovakia | 48.5 | 19.0 | ○ | | | |
| CS76910 | Hamm-1 | Sweden | 55.4 | 14.0 | ○ | | | |
| CS76914 | Haes-1 | Germany | 48.6 | 9.2 | ○ | | | |
| CS76922 | HI-4 | Germany | 48.5 | 9.0 | ○ | ○ | ○ | |
| CS76925 | Hof-1 | Germany | 48.4 | 8.9 | ○ | | | |
| CS76927 | HolA-1 2 | Sweden | 55.7 | 13.4 | ○ | | | |
| CS76931 | Hov1-10 | Sweden | 56.1 | 13.7 | ○ | | | |
| CS76932 | Hov1-7 | Sweden | 56.1 | 13.7 | ○ | ○ | | |
| CS76935 | Hov3-5 | Sweden | 56.1 | 13.7 | ○ | | | |
| CS76944 | Iasi-1 | Romania | 47.2 | 27.6 | ○ | ○ | ○ | |
| CS76956 | Jl-2 | Czech | 49.2 | 16.5 | ○ | ○ | ○ | |
| CS76959 | Kal 1 | Sweden | 56.0 | 14.0 | ○ | ○ | | |
| CS76960 | Karag-1 | Russia | 51.4 | 59.4 | ○ | ○ | | |
| CS76963 | Kardz-2 | Bulgaria | 41.7 | 25.5 | ○ | ○ | ○ | |
| CS76965 | KBG1-14 | Germany | 48.5 | 9.0 | ○ | | | |

Table S1 continued

| ABRC number | List name | Country | Latitude | Longitude | PI 5ºC | PI 10ºC | PS act. | ST |
|---|---|---|---|---|---|---|---|---|
| CS76971 | Knjas-1 | Serbia | 43.5 | 22.3 | ○ | | | |
| CS76972 | KNO1.37 | USA | 41.3 | -86.6 | ○ | ○ | ○ | |
| CS76977 | Kolyv-2 | Russia | 51.3 | 82.6 | ○ | ○ | | |
| CS76995 | IP-Lab-7 | Spain | 40.9 | -4.5 | ○ | | | |
| CS76996 | IP-Lac-0 | Spain | 43.3 | -5.9 | ○ | ○ | ○ | |
| CS77001 | Lag1-6 | Georgia | 41.8 | 46.3 | ○ | | | |
| CS77029 | Lerik2-7 | Azerbaijan | 38.8 | 48.6 | ○ | ○ | | |
| CS77030 | Leska-1-44 | Bulgaria | 41.5 | 25.0 | ○ | | | |
| CS77032 | Lesno-1 | Russia | 53.0 | 51.9 | ○ | | | |
| CS77036 | LI-OF-065 | USA | 40.8 | -72.9 | ○ | | | |
| CS77051 | IP-Loz-0 | Spain | 41.0 | -3.8 | ○ | | | |
| CS77056 | Lu-1 | Sweden | 55.7 | 13.2 | ○ | ○ | | |
| CS77057 | Lu3-30 | Germany | 48.5 | 9.1 | ○ | | | |
| CS77074 | IP-Mat-0 | Spain | 41.8 | 2.7 | ○ | ○ | | |
| CS77076 | IP-Mdd-0 | Spain | 41.9 | -2.8 | ○ | | | |
| CS77080 | Melni-2 | Bulgaria | 41.5 | 23.4 | ○ | | | |
| CS77088 | Mitterberg-3-187 | Italy | 46.4 | 11.3 | ○ | | | |
| CS77099 | MNF-Pot-21 | USA | 43.6 | -86.3 | ○ | | | |
| CS77103 | IP-Moc-11 | Spain | 41.6 | -5.6 | ○ | ○ | | |
| CS77107 | IP-Mon-5 | Spain | 38.1 | -4.4 | ○ | ○ | | |
| CS77121 | Nar-5 | Azerbaijan | 39.0 | 48.9 | ○ | | | |
| CS77128 | No-0 | Germany | 51.1 | 13.3 | ○ | | | |
| CS77137 | Nyl-7 | Sweden | 63.0 | 18.3 | ○ | | | |
| CS77156 | Oy-0 | Norway | 60.4 | 6.2 | ○ | | | |
| CS77161 | Panik-1 | Russia | 53.1 | 52.2 | ○ | | | |
| CS77165 | IP-Pdl-0 | Spain | 43.0 | -5.6 | ○ | | | |
| CS77177 | Pigna-1 | Italy | 41.2 | 14.2 | ○ | ○ | | |
| CS77178 | IP-Pil-0 | Spain | 40.5 | -4.3 | ○ | | | |
| CS77179 | IP-Piq-0 | Spain | 42.1 | -2.6 | ○ | ○ | | |
| CS77183 | PNA3.10 | USA | 42.1 | -86.3 | ○ | ○ | | |

Table S1 continued

| ABRC number | List name | Country | Latitude | Longitude | PI 5ºC | PI 10ºC | PS act. | ST |
|---|---|---|---|---|---|---|---|---|
| CS77191 | PT2.21 | USA | 41.3 | -86.7 | ○ | ○ | | |
| CS77194 | Puk-1 | Sweden | 56.2 | 14.7 | ○ | | | |
| CS77216 | IP-Ria-0 | Spain | 42.3 | 2.2 | ○ | | | |
| CS77218 | Rmx-A180 | USA | 42.0 | -86.5 | ○ | | | |
| CS77246 | Sf-1 | Spain | 41.8 | 3.0 | ○ | ○ | | |
| CS77260 | Sparta-1 | Sweden | 55.7 | 13.2 | ○ | | | |
| CS77265 | Spro 3 | Sweden | 57.3 | 18.2 | ○ | | | |
| CS77266 | Sq-1 | UK | 51.4 | -0.6 | ○ | | | |
| CS77271 | Stara-1 | Bulgaria | 42.5 | 25.6 | ○ | ○ | | |
| CS77279 | Stiav-1 | Slovakia | 48.5 | 18.9 | ○ | | | |
| CS77284 | Strand-1 | Norway | 68.8 | 15.5 | ○ | ○ | | |
| CS77301 | T510 | Sweden | 55.8 | 13.1 | ○ | | | |
| CS77306 | T580 | Sweden | 55.8 | 13.1 | ○ | ○ | | |
| CS77310 | T710 | Sweden | 55.8 | 13.3 | ○ | | | |
| CS77320 | T850 | Sweden | 55.9 | 13.6 | ○ | ○ | ○ | |
| CS77322 | T880 | Sweden | 55.9 | 13.6 | ○ | | | |
| CS77327 | T980 | Sweden | 55.9 | 13.5 | ○ | ○ | | |
| CS77342 | IP-Tau-0 | Spain | 42.5 | 0.8 | ○ | | | |
| CS77349 | TDr-17 | Sweden | 55.8 | 14.1 | ○ | | | |
| CS77352 | TDr-4 | Sweden | 55.8 | 14.1 | ○ | | | |
| CS77357 | Teano-1 | Italy | 41.3 | 14.1 | ○ | | | |
| CS77363 | TFa 07 | Sweden | 63.0 | 18.3 | ○ | ○ | | |
| CS77371 | IP-Tol-7 | Spain | 42.1 | 0.6 | ○ | | | |
| CS77373 | TOM 03 | Sweden | 63.0 | 18.4 | ○ | ○ | | |
| CS77378 | IP-Tor-1 | Spain | 41.6 | -2.8 | ○ | | | |
| CS77385 | TRE-1 | France | 48.9 | 4.1 | ○ | ○ | | |
| CS77388 | Ts-5 | Spain | 41.7 | 2.9 | ○ | | | |
| CS77392 | Tu-B2-3 | Germany | 48.5 | 9.1 | ○ | ○ | | |
| CS77397 | Tu-W1 | Germany | 48.5 | 9.0 | ○ | | | |
| CS77636 | Ale-Stenar-77-31 | Sweden | 55.4 | 14.1 | ○ | ○ | | |

Table S1 continued

| ABRC number | List name | Country | Latitude | Longitude | PI 5°C | PI 10°C | PS act. | ST |
|---|---|---|---|---|---|---|---|---|
| CS77906 | Boo2-3 | Sweden | 55.9 | 13.5 | ○ | | | |
| CS78033 | Stu-2 | Sweden | 56.5 | 16.1 | ○ | | | |
| CS78035 | T1010 | Sweden | 55.7 | 13.2 | ○ | | | |
| CS78782 | Cnt-1 | UK | 51.3 | 1.1 | ○ | | | |
| CS78783 | For-2 | UK | 56.6 | -4.1 | ○ | | | |
| CS78802 | UKSE06-362 | UK | 51.3 | 0.4 | ○ | | | |
| CS78803 | UKSE06-432 | UK | 51.2 | 0.3 | ○ | ○ | | |
| CS78817 | Ull2-3 | Sweden | 56.1 | 14.0 | ○ | ○ | | |
| CS78825 | IP-Usa-0 | Spain | 40.7 | -3.2 | ○ | | | |
| CS78829 | IP-Val-0 | Spain | 42.3 | -3.1 | ○ | | | |
| CS78840 | IP-Ven-0 | Spain | 40.8 | -4.0 | ○ | | | |
| CS78844 | IP-Vim-0 | Spain | 41.9 | -6.5 | ○ | ○ | | |
| CS78852 | Wank-2 | Germany | 48.5 | 9.1 | ○ | | | |
| CS78853 | WAR | USA | 41.7 | -71.3 | ○ | ○ | | |
| CS78857 | Ws-0.2 | Russia | 52.3 | 30.0 | ○ | | | |
| CS78860 | Xan-3 | Azerbaijan | 38.7 | 48.8 | ○ | | | |
| CS78861 | Xan-5 | Azerbaijan | 38.7 | 48.8 | ○ | ○ | | |
| CS78862 | Xan-6 | Azerbaijan | 38.7 | 48.8 | ○ | ○ | | |
| CS78866 | Yeg-5 | Armenia | 39.9 | 45.4 | ○ | ○ | | |
| CS78868 | Yeg-8 | Armenia | 39.9 | 45.4 | ○ | ○ | | |
| CS78879 | Zerev-1-35 | Bulgaria | 41.9 | 23.1 | ○ | | | |
| CS78880 | Zu-0 | Switzerland | 47.4 | 8.6 | ○ | | | |
| CS78882 | Zupan-1 | Croatia | 45.1 | 18.7 | ○ | ○ | | |
| CS78890 | IP-Cal-0 | Spain | 40.9 | -1.4 | ○ | ○ | | |
| CS78897 | Dr-0 | Germany | 51.1 | 13.7 | ○ | ○ | | |
| CS78913 | NFA-8 | UK | 51.4 | -0.6 | ○ | | | |
| CS78914 | IP-Pro-0 | Spain | 43.3 | -6.0 | ○ | ○ | | |
| CS78915 | Pt-0 | Germany | 53.5 | 10.6 | ○ | | | |
| CS78917 | Sorbo | Tajikistan | 38.4 | 68.5 | ○ | | | |
| CS78920 | Ws-2 | Russia | 52.3 | 30.0 | ○ | ○ | | |

Table S1 continued

| ABRC number | List name | Country | Latitude | Longitude | PI 5ºC | PI 10ºC | PS act. | ST |
|---|---|---|---|---|---|---|---|---|
| CS78924 | Kos-2 | Russia | 62.0 | 34.1 | ○ | | | |
| CS78940 | OOE3-1 | Austria | 48.3 | 14.7 | ○ | ○ | | |
| CS78945 | BRR23 | USA | 40.8 | -87.7 | ○ | | | |
| CS78947 | BRR60 | USA | 40.8 | -87.7 | ○ | | | |
| CS78948 | BRR107 | USA | 40.8 | -87.7 | ○ | | | |
| CS78952 | LI-RR-096 | USA | 40.9 | -72.9 | ○ | ○ | | |
| CS78953 | LI-RR-097 | USA | 40.9 | -72.9 | ○ | ○ | | |
| CS78961 | MIC-24 | USA | 41.8 | -86.4 | ○ | | | |
| CS78975 | Lak-13 | USA | 41.8 | -86.7 | ○ | ○ | | |
| CS78977 | Mdn-10 | USA | 42.1 | -86.5 | ○ | ○ | | |
| CS78979 | MNF-Pot-15 | USA | 43.6 | -86.3 | ○ | | | |
| CS78982 | MNF-Pin-40 | USA | 43.5 | -86.2 | ○ | ○ | | |
| CS78985 | MuskSP-68 | USA | 43.2 | -86.3 | ○ | ○ | | |
| CS78989 | MSGA-61 | USA | 43.3 | -86.1 | ○ | ○ | | |
| CS79000 | UKSW06-240 | UK | 50.4 | -4.9 | ○ | ○ | | |
| CS79006 | UKNW06-488 | UK | 54.4 | -2.9 | ○ | ○ | ○ | |
| CS79008 | UKID11 | UK | 57.0 | -3.4 | ○ | ○ | | |
| CS79009 | UKID71 | UK | 52.9 | -1.3 | ○ | ○ | | |
| CS79012 | CSHL-17 | USA | 40.9 | -73.5 | ○ | ○ | | |
| CS79013 | FM-10 | USA | 42.4 | -76.5 | ○ | ○ | | |
| CS79014 | FM-11 | USA | 42.4 | -76.5 | ○ | ○ | | |
| CS79016 | HS-17 | USA | 42.4 | -71.1 | ○ | ○ | | |
| CS79019 | Tol-2 | USA | 41.7 | -83.6 | ○ | ○ | ○ | |
| CS79020 | Tol-3 | USA | 41.7 | -83.6 | ○ | ○ | | |
| CS79022 | 627RMX-1MN4 | USA | 42.0 | -86.5 | ○ | ○ | | |
| CS79025 | KNO2.41 | USA | 41.3 | -86.6 | ○ | ○ | | |
| CS79027 | PT1.52 | USA | 41.3 | -86.7 | ○ | | | |
| CS79031 | LP3413.53 | USA | 41.7 | -86.9 | ○ | ○ | | |
| CS944 | An-1 | Belgium | 51.5 | 4.5 | | ○ | | |
| CS996 | Bs-1 | Switzerland | 47.0 | 7.0 | | ○ | | |

Table S1 continued

| ABRC number | List name | Country | Latitude | Longitude | PI 5°C | PI 10°C | PS act. | ST |
|---|---|---|---|---|---|---|---|---|
| CS1154 | Fe-1 | Germany | 48.0 | 8.5 | | ○ | | |
| CS1264 | Kas-2 | India | 34.0 | 74.0 | | ○ | | |
| CS1380 | Mt-0 | Libya | 33.0 | 23.0 | | ○ | | |
| CS1436 | Oy-0 | Norway | 60.0 | 6.0 | | ○ | | |
| CS1480 | Ra-0 | France | 46.0 | 3.0 | | ○ | | |
| CS1550 | Te-0 | Finland | 63.0 | 21.0 | | ○ | | |
| CS22587 | Ull-2-3 | Sweden | 55.0 | 13.0 | | ○ | | |
| CS22606 | KZ-1 | Kazakhstan | 50.0 | 59.0 | | ○ | | |
| CS22624 | Yo-0 | USA | 37.0 | -119.0 | | ○ | | |
| CS22650 | Ll-0 | Spain | 41.6 | 2.5 | | ○ | | |
| CS22678 | Br-0 | Czech | 49.0 | 16.5 | | ○ | | |
| CS22682 | Cvi-0 | Cape Verde | 15.0 | -23.0 | | ○ | | |
| CS22684 | Fei-0 | Portugal | 39.0 | -7.0 | | ○ | | |
| CS28220 | Edi-0 | UK | 56.0 | 3.0 | | ○ | | |
| CS28715 | Rsch-0 | Russia | 56.0 | 34.0 | | ○ | | |
| CS28980 | Dzi-1 | Tajikistan | 37.6 | 72.6 | | ○ | | |
| CS76423 | Galdo-1 | Italy | 40.6 | 15.3 | | ○ | ○ | |
| CS76540 | Le-0 | Netherlands | 52.2 | 4.5 | | ○ | | |
| CS76590 | Rome-1 | Italy | 42.0 | 12.1 | | ○ | | |
| CS76647 | IP-Adm-0 | Spain | 39.2 | -4.5 | | ○ | | |
| CS76770 | CIBC-17 | UK | 51.4 | -0.6 | | ○ | | |
| CS77100 | MNF-Pot-75 | USA | 43.6 | -86.3 | | ○ | | |
| CS78800 | UKSE06-252 | UK | 51.3 | 0.5 | | ○ | | |
| CS78842 | IP-Vid-1 | Portugal | 38.2 | -7.8 | | ○ | | |
| CS78908 | Mitterberg-1-180 | Italy | 46.4 | 11.3 | | | ○ | |

The information of the latitude and longitude of the habitats are gained at Arabidopsis Biological Resource Center (ABRC) and The Arabidopsis Information Resource (TAIR).

**Methods S1** Equations of energy-dependent non-photochemical quenching.

Because, in the present study, samples after photoinhibition treatment were used for the estimation of energy-dependent non-photochemical quenching, we used the following two indicators. We should also note that the photoinhibition treatment was conducted in a temperature-controllable box, where fluorometer (i.e. FluorCam) did not fit. Therefore, we measured the non-relaxed $F_v/F_m$ (i.e $F_v/F_m\mathrm{NR}$) just after the photoinhibition treatment, and measured $F_v/F_m$ after 30 min dark treatment.
In the following equations, NR means "not relaxed".

Fluorescence quantum yield of $F_o$, $F_o\mathrm{NR}$, $F_m$ and $F_m\mathrm{NR}$ can be shown as follows:
$F_o = k_F / (k_F + k_D + k_P)$,
$F_o\mathrm{NR} = k_F / (k_F + k_D + k_E + k_P)$,
$F_m = k_F / (k_F + k_D)$,
$F_m\mathrm{NR} = k_F / (k_F + k_D + k_E)$,
where $k_E$, $k_F$, $k_D$ and $k_P$ indicate rate constants for the de-excitation of excited chlorophyll molecules by energy-dependent quenching, by fluorescence, by nonradiative decay process and by photochemistry, respectively.

Therefore, the following parameters can be calculated as follows:
$F_v/F_m = (F_m - F_o)/ F_m = k_P / (k_F + k_D + k_P)$,

$F_v/F_m\mathrm{NR} = (F_m\mathrm{NR} - F_o\mathrm{NR})/ F_m\mathrm{NR} = k_P / (k_F + k_D + k_E + k_P)$,

$$F_v/F_m - (F_v/F_m)_{NR} = k_P / (k_F + k_D + k_P) - k_P / (k_F + k_D + k_E + k_P)$$
$$= k_P \cdot k_E / \{(k_F + k_D + k_P)(k_F + k_D + k_E + k_P)\}$$
$$= F_v/F_m\mathrm{NR} \cdot k_E /(k_F + k_D + k_P),$$

$qE$ indicator $= (F_v/F_m /(F_v/F_m)_{NR} - 1) = k_E /(k_F + k_D + k_P)$.

Previous studies suggested the difficulty of the measurement of $F_o\mathrm{NR}$ (as known as $F_o'$) and a calculation method is proposed by Oxborough and Baker (1997, *Photosynthesis Research* 54: 135-142). However, the necessity of the adjustment of fluorescence sensitivity in each measurement of FluorCam made it difficult to calculate $F_o\mathrm{NR}$ by the proposed method in the present research.
